# Removing Thermostat Distortions of Protein Dynamics in Constant-Temperature Molecular Dynamics Simulations

**DOI:** 10.1101/2021.06.08.447619

**Authors:** Alan Hicks, Matthew MacAinsh, Huan-Xiang Zhou

## Abstract

Molecular dynamics simulations are widely used to determine equilibrium and dynamic properties of proteins. Nearly all simulations nowadays are carried out at constant temperature, with a Langevin thermostat among the most widely used. Thermostats distort protein dynamics, but whether or how such distortions can be corrected has long been an open question. Here we show that constant-temperature simulations with a Langevin thermostat dilate protein dynamics and present a correction scheme to remove the dynamic distortions. Specifically, ns-scale time constants for overall rotation are dilated significantly but sub-ns time constants for internal motions are dilated modestly, while all motional amplitudes are unaffected. The correction scheme involves contraction of the time constants, with the contraction factor a linear function of the time constant to be corrected. The corrected dynamics of eight proteins are validated by NMR data for rotational diffusion and for backbone amide and side-chain methyl relaxation. The present work demonstrates that, even for complex systems like proteins with dynamics spanning multiple timescales, one can predict how thermostats distort protein dynamics and remove such distortions. The correction scheme will have wide applications, facilitating force-field parameterization and propelling simulations to be on par with NMR and other experimental techniques in determining dynamic properties of proteins.

**TOC graphic:** 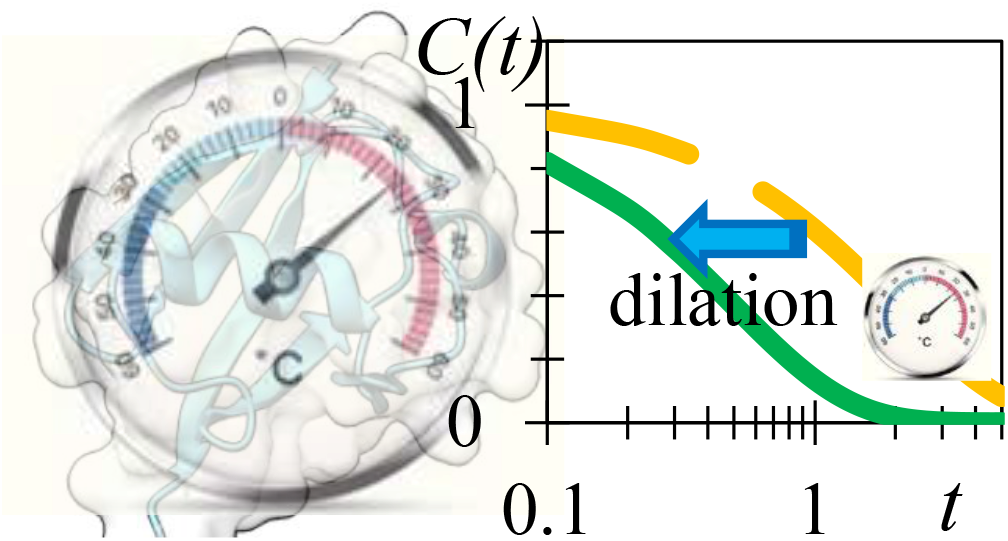

## 1. INTRODUCTION

Molecular dynamics (MD) simulations are now an indispensable tool for characterizing the thermodynamic and dynamics properties of biomolecular systems. To achieve realism and facilitate comparison with experiments, modern simulations are all done at constant temperature. A number of thermostat algorithms have been developed ^1–8^. One of the most popular choices is Langevin dynamics ^5^, where two terms are added to Newton’s equation of motion for each degree of freedom of the system:

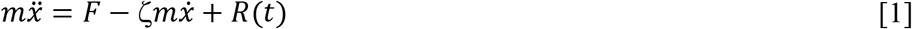

One of the added terms, *R*(*t*), is a stochastic force that thermalizes the system whereas the other, 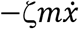, is a frictional force that damps the velocity when the temperature is too high. In the limit where the damping constant *ζ* → 0, both of the added terms vanish and one recovers the system-only equation of motion, which leads to constant energy (constant NVE) rather than constant temperature (constant NVT). Over the years, a number of artifacts of thermostats have been discovered and addressed, leading some to call thermostats a “necessary evil” ^9^. These include the “flying ice cube” problem (hot solvent-cold solute) ^10^; non-ergodicity ^6^; and distortion of configuration-space distributions ^11^. It is important to recognize that all thermostats are only designed to produce the correct equilibrium thermodynamic properties at a given temperature. All thermostats inevitably perturb the dynamic (i.e., time-dependent) properties of the biomolecular systems ^12–14^. There are only sporadic attention paid to this crucial issue ^7, 12, 15–18^. As MD simulations more and more routinely reach microseconds and beyond and direct comparison with experimental data on macromolecular dynamics become more and more practical, it is becoming pressing to assess and correct the dynamic distortions of thermostats. Here we address this problem for the Langevin thermostat.

Early studies identified the influences of a Langevin thermostat on the dynamics of small molecules. Loncharich et al. ^15^ showed that the isomerization rate of N-acetyl-methylaldehyde was dictated by the damping constant *ζ*. In line with Kramers’ theory ^19^, the isomerization rate reached a maximum at an intermediate *ζ* (around 2 ps^−1^) and became inversely proportional to *ζ* in high-friction (*ζ* > 10 ps^−1^) Langevin simulations. Similarly, the translational diffusion constants of neon and water decreased with increasing *ζ* ^16^. Tobias et al. ^12^ compared NVE and NVT simulations and found that, at a small *ζ* (= 2 ps^−1^), Langevin and Nose-Hoover thermostats did not alter the sub-ps underdamped time correlation function of the Tyr41 *χ*^2^ torsion angle in bovine pancreatic trypsin inhibitor (BPTI). However, this correlation function became overdamped in high-friction (*ζ* = 20 or 50 ps^−1^) Langevin simulations. Bussi at al. ^7^ also presented evidence of dynamic distortions when their thermostat, involving stochastic velocity rescaling, was at a high damping constant. Basconi and Shirts ^17^ investigated the effects of several thermostats on the transport dynamics of water and conformational dynamics of a coarse-grained homopolymer. They stated that, with stronger coupling to the thermal bath (e.g., by increasing the damping constant *ζ* of a Langevin thermostat), “the dynamics will begin to diverge from the NVE dynamics.” In particular, water diffusion and homopolymer chain decorrelation were both slowed down by several-fold when a Langevin thermostat at *ζ* = 10 ps^−1^ was used. Debiec et al. ^18^ reported ~50% increases in the overall rotational correlation times of three proteins (the third immunoglobulin-binding domain of protein G (GB3), ubiquitin, and binase) in Langevin simulations (at *ζ* = 1 ps^−1^) compared with NVE simulations. Recognizing the conflicting effects of thermostats in achieving thermal equilibration and producing dynamic distortions, Leimkuhler et al. ^14^ proposed using the ratio of these two effects as a measure for selecting an optimal thermostat.

Many efforts in force-field parameterization have included experimental data of dynamic properties for parameter tuning. Translational and rotational diffusion constants are often used in parameterizing water models, as illustrated by the re-parameterization of the SPC/E water model into SPC/E_b_ ^20^. NMR spin relaxation properties have long been used to validate force fields ^21–24^. In particular, Hoffmann et al. ^23^ tuned side-chain methyl rotation energy barriers in the AMBER ff99SB*-ILDN force field ^25–27^, and used NMR relaxation of methyls in ubiquitin ^28–29^ to validate calculations in NVE and NPT simulations (with Bussi at al.’s stochastic velocity rescaling ^7^ as the thermostat). This study did not reveal any significant effects of the thermostat and barostat on methyl axis order parameters and correlation times of fast internal motions. However, in theory all thermostats distort protein dynamics, and hence force-field parameters might be tuned in part to compensate for dynamic distortions of thermostats. Moreover, the extent of dynamic distortion may depend on the particular protein and on the thermostat used, therefore raising serious issues on force-field transferability.

For globular proteins, the model-free approach of Lipari and Szabo ^30^ and its extension ^31^ have been highly successful in analyzing NMR spin relaxation, for data from experiments as well as from MD simulations. This approach assumes separation between timescales of overall rotation and internal motions. For intrinsically disordered proteins, we and others carried out NVT simulations using a Langevin thermostat and borrowed the mathematical form of the model-free approach to analyze NMR relaxation ^32–34^. Specifically, the time correlation function of a backbone NH bond vector, *C*(*t*) =< *P*_2_(**n**(*t*′) ∙ **n**(*t*′ + *t*)) >_*t*′_, was fit to a sum of exponential functions of time. Here *P*_2_ is the second-order Legendre polynomial and **n** denotes the unit vector along the NH bond. In our study of ChiZ ^34^, we used three exponentials, with time constants (*τ*_*i*_; *i* = 1 to 3) at ~10 ns, ~2 ns, and ~300 ps, as well as an ultrafast (~10 ps) component. The observed patterns of the longitudinal (*R*_1_) and transverse (*R*_2_) relaxation rates along the amino-acid sequence were predicted well, but the MD simulations systematically underestimated *R*_1_ and overestimated *R*_2_, suggesting that the time constants, especially the longest one, were too long. By contracting, i.e., scaling down, the time constants by a factor 1 + *bτ*_*i*_, we significantly improved the agreement between the relaxation properties calculated from the MD simulations and the experimental data. Dynamic distortion by the Langevin thermostat was suggested as one of the reasons for the need of the contraction in *τ*_*i*_.

This study was designed to fully assess the dynamic distortions of the Langevin thermostat and develop a scheme to correct such distortions. We chose to study a set of eight globular proteins of various sizes (with the number of amino acids ranging from 18 to 129; Fig. 1). Globular proteins were chosen because their dynamics involve motions on different timescales and the nature of these motions (i.e., overall rotation and internal motions) is well-known. A wide range of protein sizes was chosen because we suspected that the extent of Langevin distortions is dependent on protein size. This suspicion originated from the time contraction factor introduced for ChiZ, which as a linear function of *τ*_*i*_ was greater for a mode with a longer time constant than for a mode with a shorter time constant; the former mode presumably involved the correlated motion of a larger group of atoms. Here dynamic distortions of the Langevin thermostat was identified by comparing against NVE simulations. The Langevin thermostat is found to dilate the time constants, but leave the motional amplitudes unaffected. Moreover, applying a contraction factor, *a* + *bτ*_*i*_, to the time constants completely removes the dynamic distortions. The corrected NVT results for NMR relaxation properties of backbone amides and side-chain methyls agree well with experimental data.

**Figure 1.**
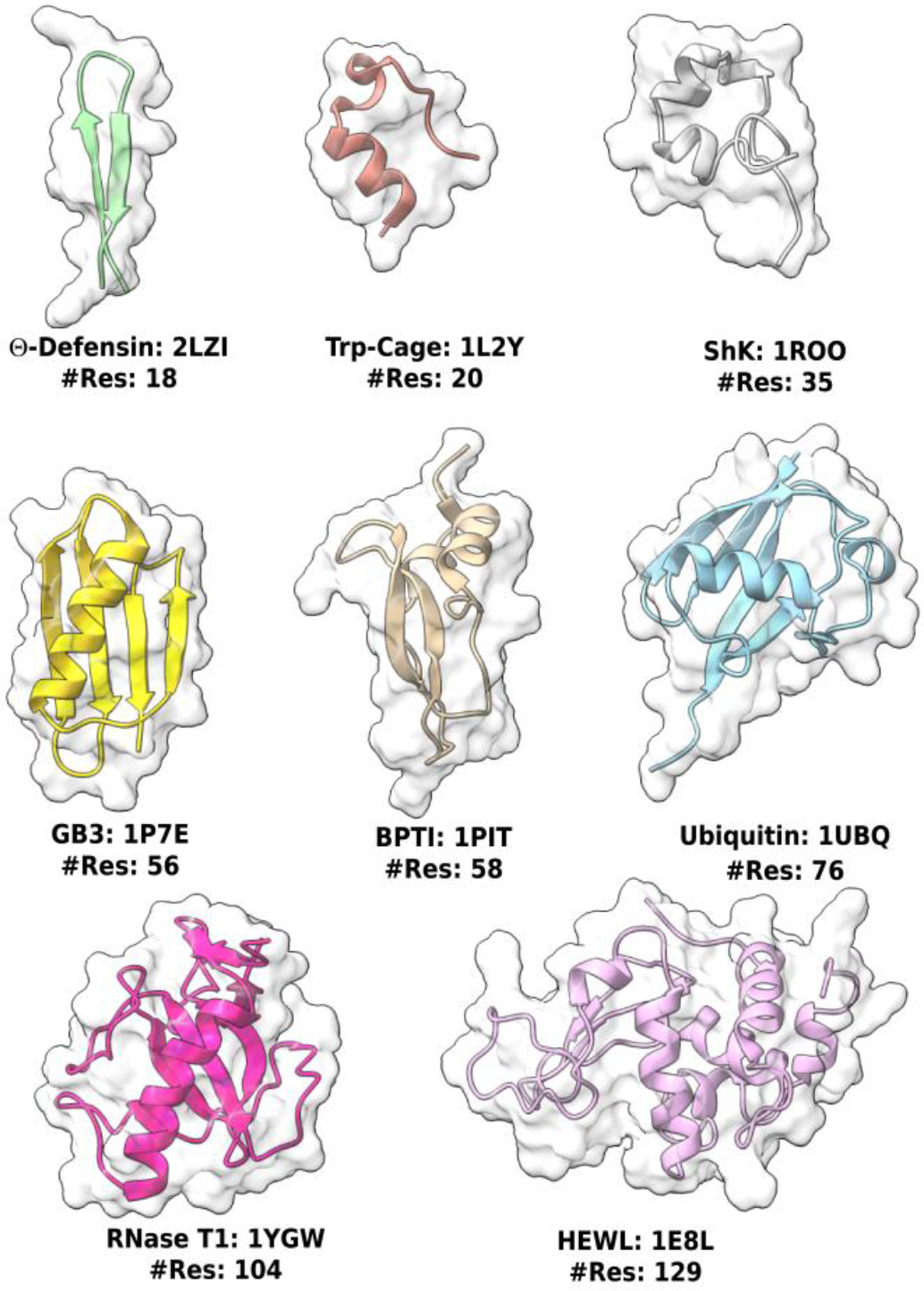
The eight proteins studied in the present work. The Protein Data Bank entry and the total number of residues of each protein are given.

## 2. COMPUTATIONAL METHODS

### 2.1. MD Simulations

Starting structures of the eight proteins were downloaded from the Protein Data Bank using the entry names listed in Fig. 1. Each structure was prepared for AMBER simulations ^35^ in *tleap* using the ff14SB protein force field ^36^ and TIP4P-D water model ^37^. The unit cell for each system was an orthorhombic box with a 15 Å solvent layer of solvent around the protein. Lastly protein-neutralizing ions (Na^+^ or Cl^−^) and NaCl (at the experimental salt concentrations) were added.

For each system, energy minimization (2000 steps of steepest descent and 3000 steps of conjugate gradient) was followed by short equilibration (100 ps of NVT and 3 ns of NPT). In the NVT portion, the temperature was ramped from 0 to the target value (Table S1) in the first 40 ps and the protein was restrained with a l kcal/mol/Å^2^ force constant. Temperature was regulated by the Langevin thermostat (damping constant at 1 ps^−1^) and pressure was regulated by the Berendsen barostat ^2^ at 1 atm. All MD simulations were carried out in AMBER18 using *pmemd.cuda* ^38^.

Four replicate simulations of each equilibrated system were run for 500 ns at constant NVE and 1000 ns at constant NVT. Snapshots were saved every 0.5 ps. Bond lengths involving hydrogens were constrained using the SHAKE algorithm ^39^. Long-range electrostatic interactions were treated by the particle mesh Ewald (PME) method ^40^. The cutoff distance for nonbonded interactions were 10 Å. The timesteps were 1 fs in the NVE simulations and 2 fs in the NVT simulations. In the latter simulations, temperature was regulated by the Langevin thermostat, with the damping constant set to each of 3 values: 0.2 ps^−1^, 2 ps^−1^, and 20 ps^−1^.

Replicates for NVE production simulations were generated by independent short NPT simulations following the equilibration stage. To control energy drift, very strict convergence levels (at 0.5 × 10^−6^) were chosen for PME and SHAKE. If the energy and temperature still drifted due to known cutoff and numerical integration artefacts in NVE simulations ^13^, then more replicate simulations were run. In most cases, at least eight NVE simulations were run and the four that had the least energy and temperature drifts were selected for analysis.

### 2.2. Secondary Structure and Solvent Accessible Surface Area (SASA)

Secondary structures and SASAs were calculated using the *dssp* and *surf* commands in *cpptraj* ^41^. The SASA for each residue was converted to the percent exposure as (1 − SASA/SASA_max_) ∙ 100%, where SASA_max_ was the nominal maximum SASA for a residue of that type (e.g., alanine).

### 2.3. Rotational Correlation Time (RCT)

RCT was calculated using the *rotdif* command in in *cpptraj* as described in Wong et al. ^13^. In brief, 500 random unit vectors attached to the protein molecular frame were used to calculate time correlation functions. The latter entailed determining the dot product between each random vector, **v**(*t*_1_), at time *t*_1_ along a simulation and the same vector, **v**(*t*_2_), at time *t*_2_, using the rotation matrices required to align the protein structures at *t*_1_ and *t*_2_ to a reference (chosen as the first saved snapshot). The second-order Legendre polynomial (i.e., *P*_2_) of this dot product as a function of the lag time *t* = *t*_2_ − *t*_1_ was averaged (over the simulation and the random vector) and integrated to yield the RCT for isotropic tumbling. Finally, an average of RCTs was taken over the four replicate simulations.

### 2.4. NMR Spin Relaxation Properties

The backbone ^1^H-^15^N NMR relaxation properties were calculated from the NH bond correlation function, *C*(*t*) =< *P*_2_(**n**(*t*′) ∙ **n**(*t*′ + *t*)) >_*t*′_. *C*(*t*) was obtained using the *timecorr* function in *cpptraj*. After averaging over the four replicate simulations, it was fit to a bi-exponential function,

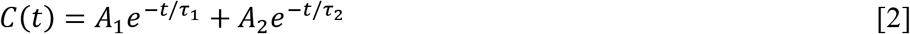

The longer time constant *τ*_1_ is for overall rotation whereas the shorter time constant *τ*_2_ is for internal motion. We did not treat *τ*_1_ as a global parameter but made it residue-specific, just as *τ*_2_ was residue-specific. Nor did we constrain the sum of the amplitudes, *A*_1_ + *A*_2_, to 1. The fitting was performed using the trust region reflexive algorithm in the *scipy.optimize.curve_fit* module of python, with *τ*_2_ bounded between 0.5 ps and 150 ps. The time constants *τ*_1_ and *τ*_2_ (as do the amplitudes *A*_1_ and *A*_2_) have large disparities. To ensure a proper balance between the portion of the data that determined the first exponential and the portion of the data that determined the second exponential, in the fitting we picked data points that were uniformly distributed on a log scale of time instead of a linear scale of time. Specifically, we used only data points with indices given by ⌈1.02^*i*^⌉, where ⌈*x*⌉ denotes the ceiling function (i.e., the smallest integer greater than or equal to *x*) and *i* = 0, 1, 2, …

From the bi-exponential fit, the spectral density was found as

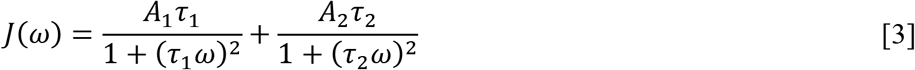

Finally *R*_1_, *R*_2_, and NOE were given by

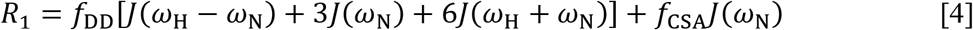

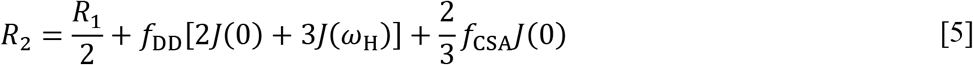

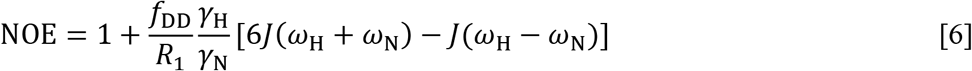

where 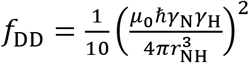 and 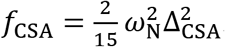. The meanings of the symbols are: *μ*_0_, permittivity of free space; ℏ, reduced Plank constant; γ_N_ and γ_N_, gyromagnetic ratios of nitrogen and hydrogen; *ω*_H_ = γ_N_*B*_0_, Larmor frequency of hydrogen; *ω*_N_, counterpart of nitrogen; *r*_NH_, NH bond length (set at 1.02 Å); and Δ_CSA_ (= −170 ppm), chemical shift anisotropy of nitrogen.

The *A*_1_ amplitude of the bi-exponential fit can be directly interpreted as the order parameter *S*^2^ (the A1 method). Order parameters can also be determined after removing overall rotation by structural alignment. Alignment was done with each block of an MD trajectory. The block length should be between *τ*_1_ and *τ*_2_, and was chosen to be 1 ns for GB3 and ubiquitin. Longer block lengths were also tested. After alignment with the first snapshot of the block, the order parameter was calculated as an average second-order Legendre polynomial ^42^:

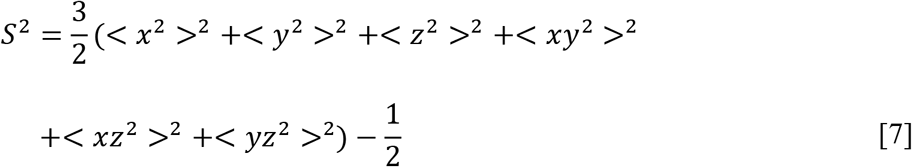

where (*x*, *y*, *z*) denote the Cartesian components of the unit vector along an NH bond, and < ⋯ > signifies averaging over the block. A final average of the *S*^2^ values from the different blocks was taken. We call the foregoing procedure the P2 method.

Relaxation properties for side-chain methyls were calculated in much the same way. The CH correlation functions were averaged over the three hydrogens in the methyl, and fit to a bi-exponential function (with *τ*_2_ bounded between 0.5 ps and 350 ps) to produce the spectral densities. The in-phase transverse and longitudinal relaxation rates were finally calculated as

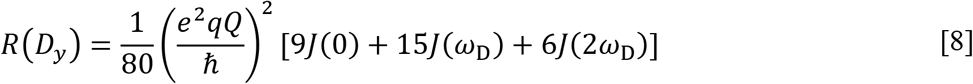

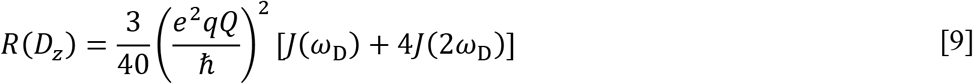

with the quadrupolar coupling constant 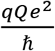 set to 167∙ 2*π* kHz ^43^ and *ω*_D_ denoting the Larmor frequency of deuterium. Methyl axis order parameters, 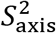, were calculated from CC correlation functions (A1 method).

### 2.5. Time-contraction scheme

The time-contraction factor was obtained as the ratio between the RCT of a set of replicate NVT simulations and the NVE counterpart (Eq [10]). A linear correlation analysis (Eq [11]) was then carried out in Excel between this factor and RCT_NVT_, using pairs of such values for the eight proteins studied. Values of the *a* and *b* coefficients were obtained for the Langevin thermostat at three damping constants (0.2 ps^−1^, 2 ps^−1^, and 20 ps^−1^), and then used to parameterize *a* and *b* as functions of the damping constant *ζ* (Eqs [13] and [14]).

## 3. RESULTS

For each of the eight proteins displayed in Fig. 1, we carried out four replicate NVE simulations and four replicate NVT simulations each with the Langevin thermostat at *ζ* = 0.2, 2, 20 ps^−1^. The latter simulations are referred to as NVT – 0.2, NVT – 2, and NVT – 20, respectively. We analyzed three types of dynamic properties: rotational correlation times; backbone NH time correlation functions; and side-chain methyl time correlation functions. We used the NVE simulations as benchmarks to identify dynamic distortions of the Langevin thermostat and to find a correction scheme. The corrected NVT results are validated by experimental data.

### 3.1. Increase in RCTs by Langevin Thermostat, and a Tentative Correction

In Fig. 2A, we compare the RCTs from the NVE and NVT – 2 simulations against the experimental data for the eight proteins ^44–51^ (see Supporting Table S1). As expected, RCTs increase with protein size, for both the experimental data and for the NVE and NVT results. For example, the NVE results increase from ~1 ns for the two smallest proteins (θ-defensin and Trp-cage) to ~6 ns for the largest protein (hen egg-white lysozyme (HEWL)). These NVE results closely match the experimental values; a linear correlation analysis yielded a slope of 1.01 and an *R*^2^ of 0.988 (not shown), both very close to an ideal value of 1. In contrast, the NVT – 2 simulations consistently overestimate the RCTs, with the extent of overestimating growing with protein size. Indeed, the NVT – 2 results also show a strong correlation with the experimental values (*R*^2^ = 0.989), but with an exaggerated slope of 2.08 (Fig. 2A, green line).

**Figure 2.**
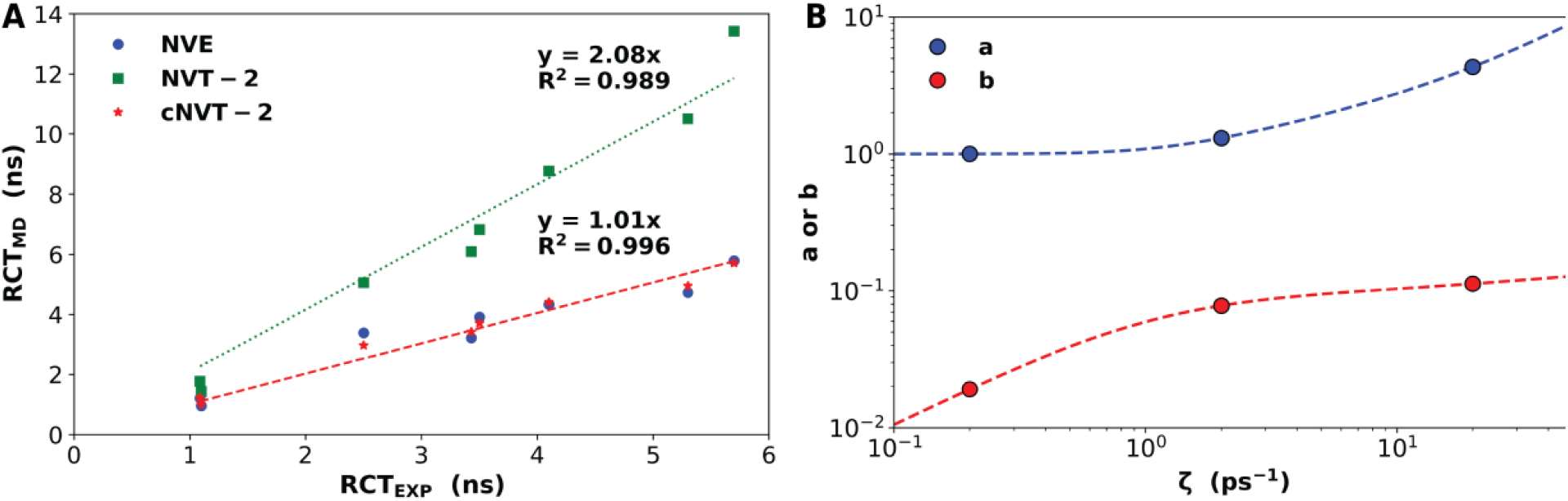
Rotational correlation times (RCTs) and scaling coefficients. (*A*) RCTs from NVE simulations show excellent agreement with experimental values, whereas those from NVT simulations (at damping constant *ζ* = 2 ps^−1^) are overestimated but, among the eight proteins, show strong correlation with the experimental values. The overestimation is completely removed after contraction by a factor *α* = *a* + *b* ∙ RCT_NVT_. The coefficients *a* and *b* were obtained taking RCT_NVT_/RCT_NVE_ as the contraction factor and doing a correlation analysis between it and RCT_NVT_ among the eight proteins. (*B*) Dependences on *a* and *b* on the damping constant *ζ* of the Langevin thermostat. Symbols are from correlation analysis; curves are functions, given by Eqs [13] and [14], designed to pass through the symbols. The values of the constants are *α*_1_ = 0.09; *β*_1_ = 1.11; *γ*_1_ = 2.01; *α*_2_ = 0.059; *β*_2_ = 0.19; and *γ*_2_ = 0.62.

It is clear that the Langevin thermostat dilates RCTs. A simple way to correct the dynamic distortion is to contract RCTs (and other time constants) calculated from Langevin simulations. We hypothesized that the ratio between the RCT calculated from NVT simulations and the NVE counterpart:

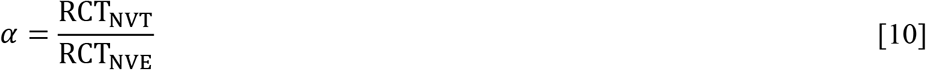

could serve as the time-contraction factor. Interestingly, as we suspected, this correction factor shows a growing trend with increasing RCT_NVT_ (or protein size; Fig. S1). Analysis based on a linear correlation,

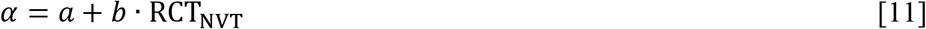

yielded *a* = 1.31 and *b* = 0.078 ns^−1^ for a Langevin thermostat at *ζ* = 2 ps^−1^. For any time constant *τ*(NVT) (including RCTs) calculated from NVT simulations, we apply a time-contraction factor *a* + *bτ*(NVT), leading to the corrected NVT (cNVT for short) time constant

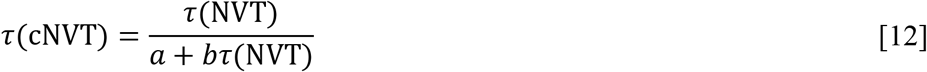

Applying this correction to the RCTs of the NVT – 2 simulations produces results that are in excellent agreement with the experimental values. A linear correlation analysis yielded a slope of 1.01 and an *R*^2^ of 0.996 (Fig. 2A, star symbols and red line).

To develop Eq [12] into a universal correction scheme for an arbitrary damping constant, we carried out additional Langevin simulations at *ζ* = 0.2 and 20 ps^−1^. The coefficients of the linear correlation in Eq [11] were found to be *a* = 1.001 and *b* = 0.019 ns^−1^ at *ζ* = 0.2 ps^−1^ and *a* = 4.33 and *b* = 0.113 ns^−1^ at *ζ* = 20 ps^−1^. The *a* and *b* values at the three *ζ* values were used to parameterize the following functional representations (Fig. 2B):

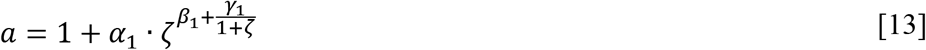

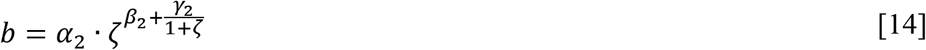

By design, these functions have the appropriate values, 1 for *a* and 0 for *b*, at *ζ* = 0 (equivalent to constant NVE). Both the *a* and *b* functions then gradually increase with increasing *ζ*. We expect that these functions, when used in Eq [12], provide the necessary time contraction for the Langevin thermostat with *ζ* in the range from 0.1 to 50 ps^−1^, which covers almost all the values of the damping constant found in simulations.

### 3.2. Langevin Thermostat Affects the Time Constants but Not Amplitudes of Backbone NH Correlation Functions

The time correlation function of a backbone NH bond involves protein dynamics on timescales ranging from ps to at least ns, and can be detected by NMR relaxation experiments. From the NVE and NVT simulations, we calculated the correlation functions of all the NH bonds in each of the eight proteins, and fit them to a sum of two exponentials (Eq [2]). NH correlation functions of three residues from NVT – 2 simulations of ubiquitin, along with the bi-exponential fits, are illustrated in Fig. S2. The amplitudes (*A*_1_ and *A*_2_) and time constants (*τ*_1_ and *τ*_2_, with *τ*_1_ > *τ*_2_) of the fits, along with secondary structure and solvent percent exposure of each residue, are displayed in Fig. 3 for GB3 and ubiquitin and in Fig. S3 for the other six proteins.

**Figure 3.**
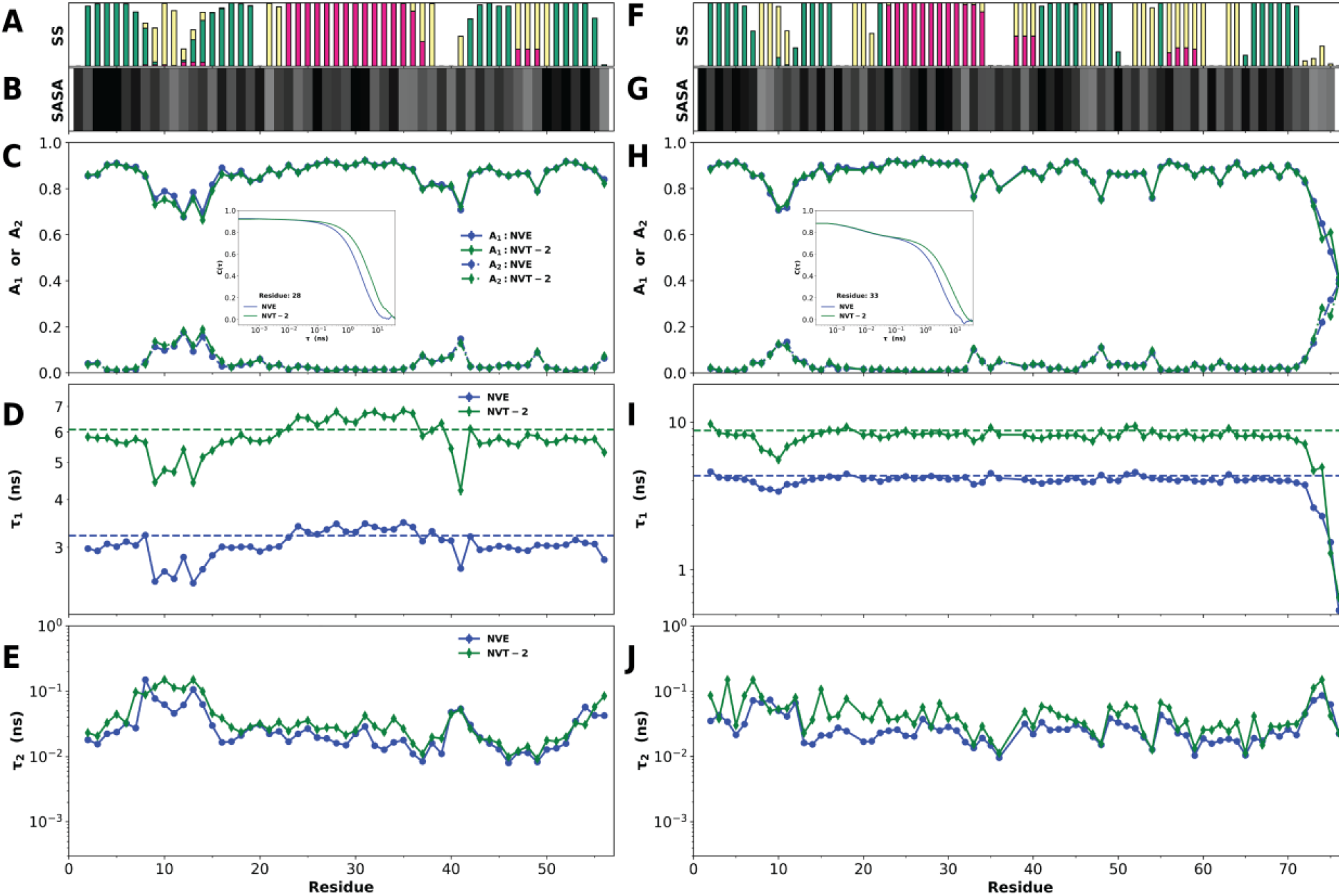
Parameters of bi-exponential fits for NH time correlation functions. (*A*-*E*) are for GB3 and (*F*-*J*) are for ubiquitin. Results for NVE and NVT (at *ζ* = 2 ps^−1^) simulations are shown. (*A*) & (*F*) Secondary structure (SS), according to the following color scheme: magenta, helix; green, β-sheet, yellow, turn; and white, coil. SS was assigned to the type with the highest propensity. (*B*) & (*H*) Percent of solvent exposure of each residue presented on a gray scale (black, buried; white, exposed). (*C*) & (*H*) Amplitudes *A*_1_ and *A*_2_ of the bi-exponential fits. Insets display the correlation functions for residue 28 in GB3 and residue 33 in ubiquitin. (*D*) & (*I*) Time constant *τ*_1_. The rotational correlation times calculated from these simulations are displayed as dashed horizontal lines. (*E*) & (*J*) Time constant *τ*_2_.

The NVE results can be summarized as follows. Most *A*_1_ values are in the 0.8 to 0.9 range, whereas most *A*_2_ values are in the 0 to 0.1 range. The sum *A*_1_ + *A*_2_ is fairly constant at around 0.9, with the “missing” amplitude (~0.1) attributable to an ultrafast (sub-ps) component. For most residues, *τ*_1_ values are close to the RCTs (1 to 6 ns) of the respective proteins, but can be much lower for flexible loops, e.g., residues 9-14 in GB3 (Figs. 3D and 1), or flexible termini, e.g., the C-terminus of ubiquitin (Figs. 3I and 1). These flexible residues, also distinguished by lower-than-average *A*_1_ (and higher-than-average *A*_2_), are outside α-helices and β-sheets and also solvent-exposed. Most *τ*_2_ values fall in the 10-100 ps range, and do not seem to be correlated with secondary structure or solvent exposure.

Comparing these NVE fitting parameters with the NVT – 2 counterparts, a striking observation is that the Langevin thermostat does not affect the amplitudes. The NVE and NVT – 2 simulations are totally independent and yet, residue by residue, *A*_1_ (*A*_2_) values of the two types of simulations track each other extremely well. The most noticeable effect of the Langevin thermostat is on *τ*_1_ values, which suffer a dilation similar to that described above for RCTs. Still, the Langevin thermostat preserves the profile of the *τ*_1_ trace over residue number, as if suggesting a constant ratio between NVT – 2 and NVE *τ*_1_ values among all the residues of a protein. For most residues, the Langevin thermostat also increases *τ*_2_ values, though to a much smaller extent than for *τ*_1_.

The forgoing effects of the Langevin thermostat – great on *τ*_1_, moderate on *τ*_2_, and null on *A*_1_ and *A*_2_ – can also be detected by directly comparing NH correlation functions from NVE and NVT – 2 simulations. Residue 28 in GB3 represents a rigid residue, located in middle of the central helix (residues 23-36; Fig. 1). As shown in Fig. 3C inset, the NH correlation functions of this residue are essentially a single exponential in both NVE and NVT – 2 simulations, but the time constant, on the ns scale, is clearly increased in the latter simulations, as indicated by a significant rightward shift of the rapidly decaying part of the correlation function (on a log-linear plot). Consistent with this qualitative difference, the bi-exponential fits yielded *A*_1_ = 0.91 and a very small *A*_2_ = 0.01 for both the NVE and NVT – 2 simulations, but with *τ*_1_ values, at 3.45 ns and 6.78 ns, differing by nearly a factor of 2. These *τ*_1_ values track closely the corresponding RCTs (3.22 ns and 6.09 ns) of GB3. On the other hand, residue 33 in ubiquitin represents a more flexible residue, located at the end of the major helix (Fig. 1). The NH correlation functions of this residue (Fig. 3H inset) show a clear bi-exponential decay. The first decays (on the 10 ps timescale) of the NVE and NVT – 2 correlation functions track each other closely. However, the second decays (on the ns timescale) deviate from each other, with the NVT – 2 time constant again about 2-fold longer than the NVE counterpart.

### 3.3. Dilation of *τ*_1_ and *τ*_2_ by Langevin Thermostat Grows with Increasing *ζ*

The benchmarking in backbone dynamics against NVE simulations has so far demonstrated that the Langevin thermostat at *ζ* = 2 ps^−1^ dilates *τ*_1_ significantly and *τ*_2_ moderately, but leaves the amplitudes of the global and local motions intact. To attain a complete picture on the effects of the Langevin thermostat, we now report on the results of NVT – 0.2 and NVT – 20 simulations.

As illustrated by GB3 and ubiquitin (Fig. S4), as the damping constant expands to a wide range (from 0.2 to 20 ps^−1^), the *A*_1_ and *A*_2_ amplitudes remain unaffected by the Langevin thermostat. Mirroring the results for RCTs (Fig. 2B and Table S1), the dilation in *τ*_1_ is smaller at *ζ* = 0.2 ps^−1^ than at *ζ* = 2 ps^−1^, but reaches 10-fold at *ζ* = 20 ps^−1^. As already suggested by results at *ζ* = 2 ps^−1^, the Langevin thermostat preserves the profile of the *τ*_1_ trace over residue number. As for *τ*_2_, at *ζ* = 20 ps^−1^ there is clearly dilation for all but one or two residues in each protein, with the *τ*_2_ profile over residue number also largely preserved.

In Figs. 4 and S5, we plot *τ*_1_ and *τ*_2_ values of each protein as violin plots, one each for NVE, NVT – 0.2, NVT – 2, and NVT – 20 simulations. Each violin plot displays the distribution of *τ*_1_ or *τ*_2_ values for all the residues of a protein, as well as the quartiles and the mean value. The relatively short heights of the *τ*_1_ violins, spanning a small fraction of a decade, reflect the fact that most *τ*_1_ values are close to the RCTs of the respective proteins. An exception is the outlying tail in Fig. 4B, due to the very flexible C-terminal residue in ubiquitin. For each protein, the mean *τ*_1_ values increase monotonically from NVE to NVT – 20.

**Figure 4.**
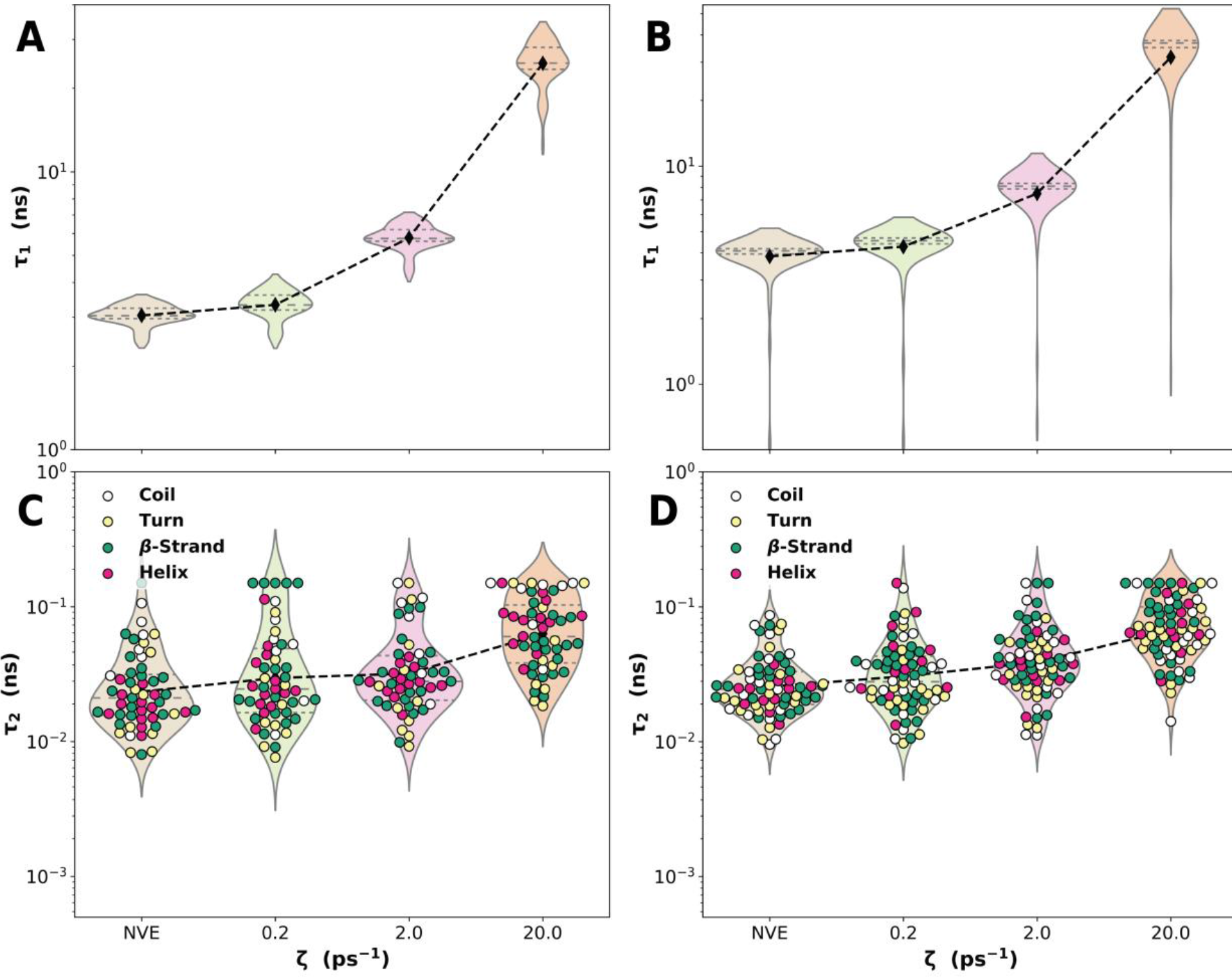
Dependences of the *τ*_1_ and *τ*_2_ time constants on the damping constant of the Langevin thermostat. NVE simulations correspond to *ζ* = 0. (*A*) & (*C*) are for GB3 and (*B*) & (*D*) are for ubiquitin. (*A*) & (*B*) *τ*_1_ values for each protein at a given *ζ* are displayed as a violin plot. Quartiles are indicated by two horizontal dashed lines in each violin plot. The mean *τ*_1_ of the protein is shown as a black diamond; these mean values at different *ζ* are connected by dashed lines between violin plots. The outlying tail in (*B*) arises from the excessively low *τ*_1_ of the very C-terminal residue in ubiquitin. (*C*) & (*D*) Corresponding data for *τ*_2_. In addition, *τ*_2_ values for individual residues are shown as a scatter plot, colored according to secondary structure.

The *τ*_2_ violins, on the other hand, cover the entire decade from 10 to 100 ps and often go beyond in both directions. As already noted, *τ*_2_ values do not seem to be correlated with secondary structure or solvent exposure. Here we further explore this matter by displaying the individual *τ*_2_ values as a scatter plot, colored according to secondary structure. For the most part, secondary structures indeed appear randomly distributed in the scatter plots. An exception occurs in a few *τ*_2_ violin plots with outlying tails, for Trp-cage, ribonuclease (RNase) T1, and HEWL in NVE and NVT – 0.2 simulations. These outlying tails originate from rigid residues located in α-helices or β-sheets. The fact that these *τ*_2_ outliers occur in NVE and NVT – 0.2 simulations but not when the Langevin thermostat is applied with the damping constant at ≥ 2 ps^−1^ suggest that rigid residues located in α-helices or β-sheets residues may be prone to being trapped in local minima in NVE and NVT – 0.2 simulations (see below).

Given the wide range of *τ*_2_ values of each protein in a given set of simulations, the trend of mean *τ*_2_ values on going from NVE simulation to NVT simulations with increasing damping constants is less clear-cut compared with the mean-*τ*_1_ trend. The change in mean *τ*_2_ could even be influenced by a few outliers (e.g., for ShK on going from NVE to NVT – 0.2). Still, there definitely is an overall increasing trend of mean *τ*_2_ going from NVE to NVT – 20.

### 3.4. Correction for *τ*_1_ and *τ*_2_ Dilated by Langevin Thermostat

As concluded above, *τ*_1_ traces from NVE and NVT simulations have similar profiles over residue number, and typical *τ*_1_ values track the corresponding RCTs. It thus seems obvious that we should apply Eq [12] to correct the dilation of *τ*1 by the Langevin thermostat, where the *a* and *b* coefficients in the time contraction factor are determined by comparing RCTs of NVE and NVT simulations. In Fig. 5A we compare the thus corrected *τ*1 for NVT – 2 (referred to as cNVT – 2) against the NVE benchmark. They show a strong correlation, with a near ideal slope of 1.01 and a very high *R*^2^ of 0.996.

**Figure 5.**
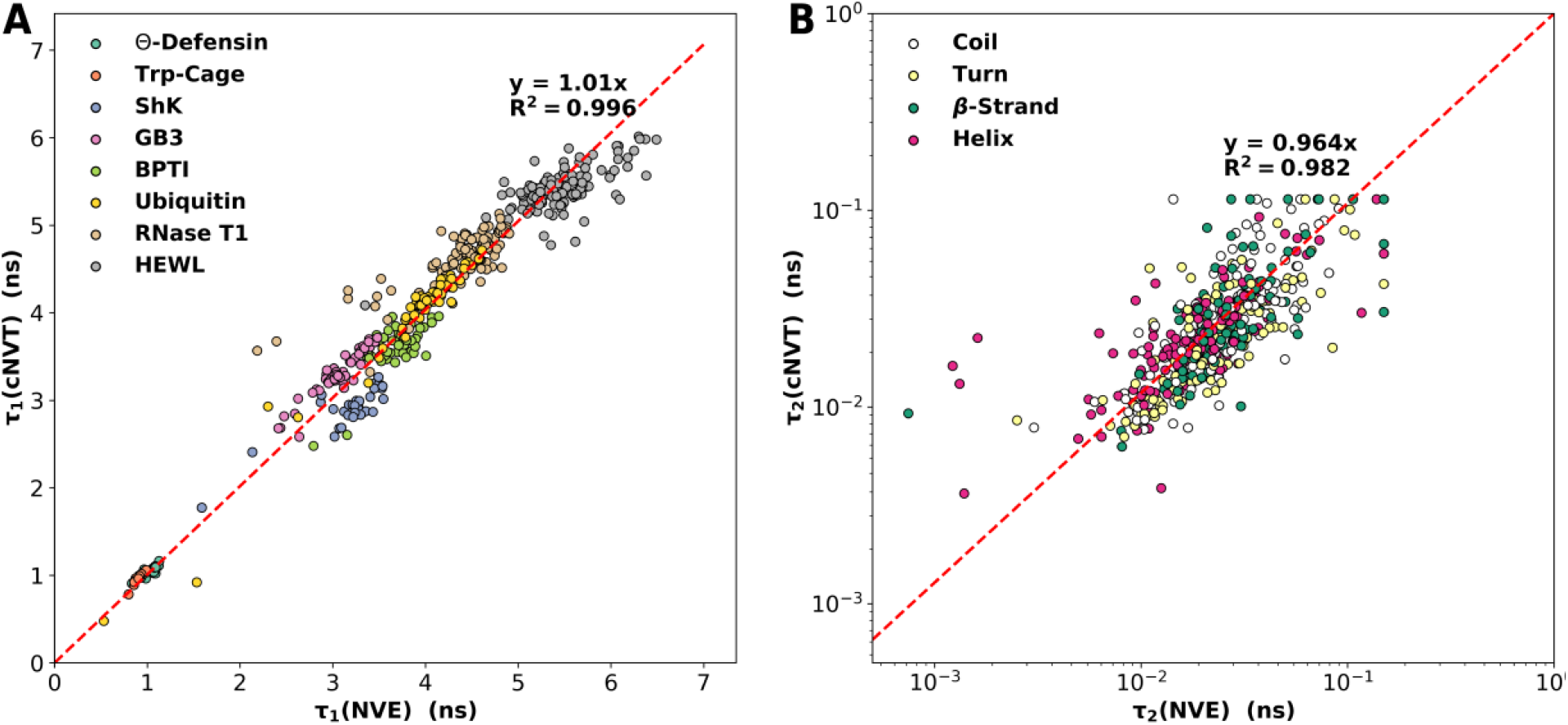
Comparison of corrected NVT time constants against the NVE results. (*A*) *τ*_1_ values for all the residues in the eight proteins from NVT simulations (at *ζ* = 2 ps^−1^), after correction by time contraction, and the NVE results are compared on a scatter plot. The correlation line and *R*^2^ value are also shown. (*B*) Corresponding data for *τ*_2_, except the correlation analysis was done on a log instead of linear scale.

Encouraged by the excellent outcome of the *τ*_1_ correction, we applied Eq [12], with the same *a* and *b* coefficients, to contract *τ*_2_. The resulting *τ*_2_ is compared with the NVE benchmark in Fig. 5B. The agreement is also good, with a linear correlation showing a slope of 0.964 and *R*^2^ of 0.982. Because the Langevin thermostat does not affect the *A*_1_ and *A*_2_ amplitudes, we expect NVT simulations after the time-constant contraction to represent the actual protein dynamics. Next we turn to experimental NMR relaxation data for validating cNVT.

### 3.5. cNVT Accurately Predicts Backbone NMR Relaxation Properties

The NH time correlation function determines the longitudinal (*R*_1_) and transverse (*R*_2_) relaxation rates of the ^15^N spin and the ^1^H-^15^N heteronuclear Overhauser effect (het. NOE). *R*_1_ and *R*_2_ are sensitive to ns dynamics and hence to *A*_1_ and *τ*_1_, whereas NOE is sensitive to both ns dynamics and sub-ns dynamics (i.e., *A*_2_ and *τ*_2_).

As illustrated in Fig. S6A for GB3 and Fig. S6B for ubiquitin, *R*_1_, *R*_2_, and NOE calculated from NVE simulations are in very good agreement with experimental values ^47, 49^. The calculated *R*_2_ values for GB3 have a root-mean-square-error (RMSE) of 0.33 s^−1^, compared to the mean *R*_2_ value of approximately 5 s^−1^. Both the low *R*_2_ values of the flexible 9-14 loop and the high *R*_2_ values of the rigid central helix (residues 23-36) are correctly predicted. NOE is also well predicted, with an RMSE of only 0.04 (compared to the mean NOE that is between 0.7 and 0.8 and typical of rigid globular proteins). The RMSE for *R*_1_ is 0.18 s^−1^, due to small but systematic overestimation. For ubiquitin, the RMSEs are 0.55 s^−1^ for *R*_2_, 0.17 s^−1^ for *R*_1_, and 0.10 for NOE, with each property showing slight overestimation.

However, *R*_2_ is significantly overestimated while *R*_1_ is significantly underestimated in NVT – 2 simulations (see Fig. S6C for GB3 and Fig. S6D for ubiquitin). The over- and underestimation get much worse in NVT – 20 simulations (Fig. S6E&F). These are clear signs that *τ*_1_ is too long. NOE overestimation is also observed in NVT – 2 simulations (Fig. S6C&D), and can largely be attributed to *τ*_1_ dilation by the Langevin thermostat. In NVT – 20 simulations, NOE overestimation does not get worse but rather is moderated; for the more flexible residues (e.g., the 9-14 loop in GB3) NOE actually becomes too low (Fig. S6E&F). The change in NOE behavior from NVT – 2 to NVT – 20 simulations is due to the fact that NOE has a non-monotonic dependence on *τ*_1_ and on *τ*_2_ ^31^. NOE increases when *τ*_1_ is dilated by 2-fold as in NVT – 2 simulations, but starts to decrease upon further *τ*_1_ dilation. The decrease is exacerbated by *τ*_2_ dilation, and becomes excessive for flexible residues with higher *A*_2_, leading to the changed NOE behavior in NVT – 20 simulations.

These problems caused by the Langevin thermostat are eliminated when the time contraction of Eq [12] is applied to *τ*_1_ and *τ*_2_. In Figs. 6 and S7 we compare cNVT – 2 calculations of *R*_1_, *R*_2_, and NOE for the eight proteins against the experimental data. These corrected NVT results have essentially the same high accuracy as the NVE results. Moreover, the correction works well for NVT simulations at the high damping constant of 20 ps^−1^ (Fig. S6G&H). For the eight proteins, NVE, cNVT – 0.2, cNVT – 2, and cNVT – 20 all have relatively small and comparable RMSEs in *R*_1_, *R*_2_, and NOE (Table S2). All the calculations miss high *R*_2_ values attributable to exchange between different chemical environments, including residues 4, 5, 7, and 11 in Trp-cage ^45^, residues 9, 18, 19, 21, 25, 31, 33, and 34 in ShK ^46^, and residues 14, 15, and 39 in BPTI ^48^. Also, at 300 K, Trp-cage undergoes partial unfolding ^45^, which probably explains why the experimental *R*_2_ values are overall higher than the calculated ones (Fig. S7B). These misses are expected, since exchange and partial unfolding processes occur on timescales longer than our MD simulations.

**Figure 6.**
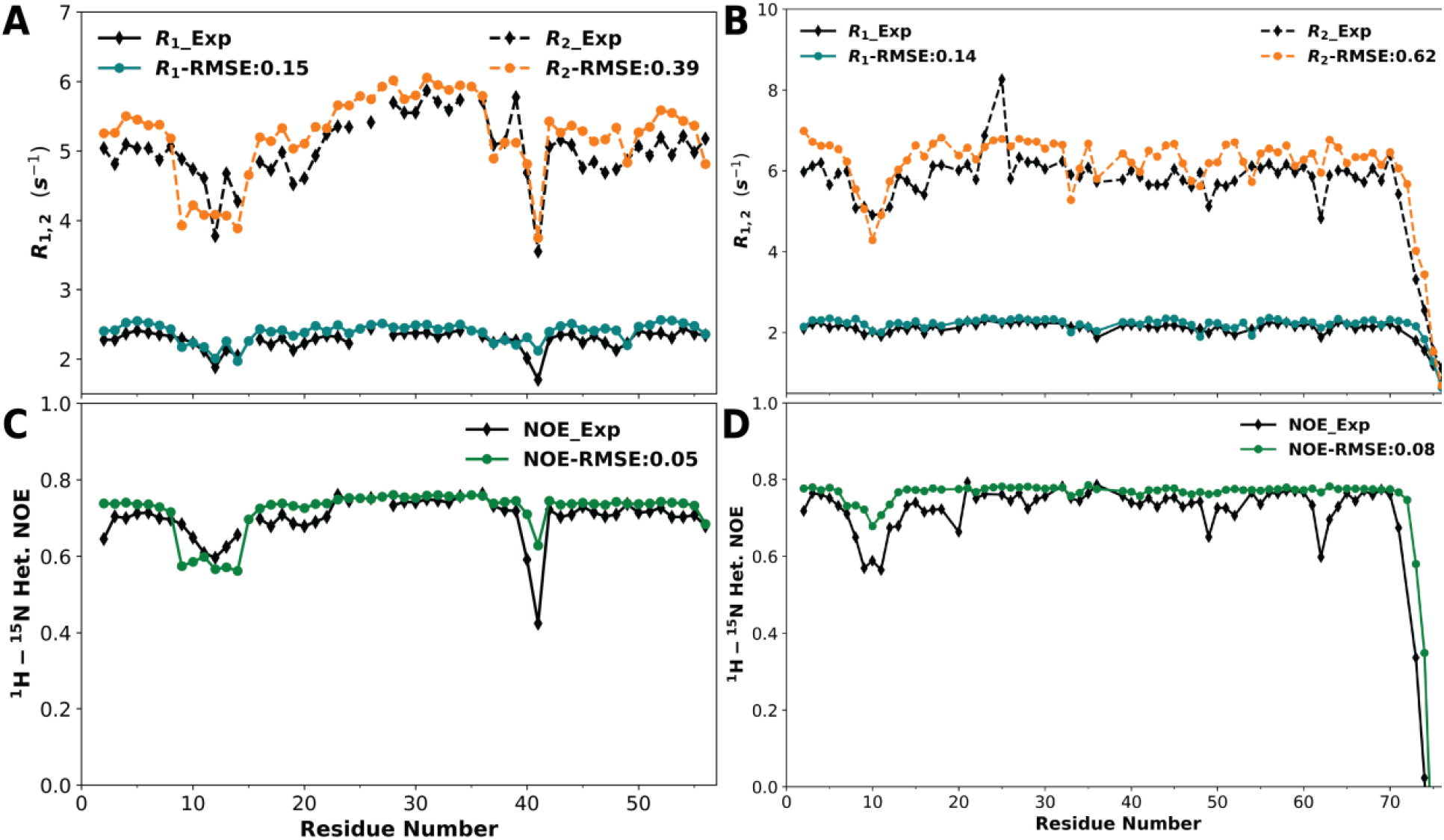
Comparison of backbone NH NMR relaxation properties predicted by corrected NVT simulations against experimental data. (*A*) & (*C*) are for GB3 and (*B*) & (*D*) are for ubiquitin. (*A*) & (*B*) Longitudinal and transverse relaxation rates from experiment (black) and from corrected NVT (at *ζ* = 2 ps^−1^; color). The root-mean-square-error (RMSE) calculated among all the residues of each protein is shown in the legend. (*C*) & (*D*) Corresponding comparison for ^1^H-^15^N heteronuclear Overhauser effect.

The spectral density of our bi-exponential fit (Eq [3]) has the same mathematical form as that in the extended model-free approach (given by Eq [4] of Clore et al. ^31^), with our *A*_1_ playing the role of the order parameter *S*^2^. In both formulations, the amplitude sum (*A*_1_ + *A*_2_) is not constrained to 1, with the deficit from 1 implying an ultrafast component. One distinction is that we treated *τ*_1_ as residue-specific but Clore et al. treated *τ*_1_ as a global parameter for overall rotation. Traditionally, order parameters have been calculated as an average second-order Legendre polynomial (*P*_2_) after removing overall rotation by structural alignment ^42^. We found that *A*_1_ matches with the average *P*_2_ if structural alignment is done within blocks of a length that is shorter than *τ*_1_ but longer than *τ*_2_, such that the average in *P*_2_ captures conformational sampling due to sub-ns internal motions (Fig. S8A&B for GB3 and ubiquitin, with 1-ns blocks for structural alignment). Indeed, *A*_1_ represents the residual order after the sub-ns decay (modeled by time constant *τ*_2_). In Fig. S8C&D, we compare order parameters calculated using the P2 method with 1-ns blocks from NVE and NVT simulations, against each other and also against experimental data ^47, 49^. This comparison confirms once again that the Langevin thermostat does not affect motional amplitudes. Compared with the experimental data, the calculated order parameters show slight overestimation overall and larger overestimation for a few of the more flexible residues (e.g., residue 12 in GB3). It is possible that the experimental *S*^2^ has contributions from internal motions on timescales longer than 1 ns. When we increased the length of the blocks for averaging, the calculated order parameters did get reduced, but the reduction could be excessive for flexible residues (not shown).

### 3.6. cNVT also Predicts Well Side-Chain NMR Relaxation Properties

To verify that our findings on the distortions of the Langevin thermostat and our correction scheme are not limited to backbone dynamics, we also calculated side-chain methyl CH and CC correlation functions from the NVE and NVT simulations. These calculated results can be directly tested by experimental data on methyl relaxation.

As illustration, in Fig. S2 we compare the CH correlation functions of three methyl-containing residues against the backbone NH correlation functions of the same residues, all from NVT – 2 simulations of ubiquitin. A clear distinction is that NH bonds undergo limited wobbling so that the sub-ns decay has a very small amplitude (small *A*_2_ and large *A*_1_ as noted above), while CH wobbling spans nearly an entire hemisphere on top of the preceding CC bonds, leading to a large amplitude for the sub-ns decay and a small amplitude for the ns decay. Accordingly, the value ranges, 0 to 0.1 for *A*_1_ and 0.6 to 0.8 for *A*_2_ from bi-exponential fits for CH bonds are the reverse of those for NH bonds (Fig. S9A). This contrast makes side-chain methyls a good test ground for our understanding of the Langevin thermostat. Despite the reversal in magnitudes for the two amplitudes (*A*_1_ and *A*_2_), the Langevin thermostat still does not perturb either amplitude. Moreover, the effects of the Langevin thermostat on the two time constants (*τ*_1_ and *τ*_2_) of CH bonds are also similar to those observed for backbone NH bonds, i.e., significant dilation on *τ*_1_ and slight change in *τ*_2_ (Fig. S9B&C). The *τ*_1_ values of CH bonds in both NVE and NVT – 2 simulations are still comparable to the respective RCTs, but have much greater site-to-site variations than found for backbone NH bonds (compare Figs. 3I and S9B). The greater variations likely reflect the fact that methyls are less tightly coupled to the protein matrix than backbone amides.

After applying the same correction scheme (Eq [12]) to *τ*_1_ and *τ*_2_ of CH bonds from NVT simulations, we calculated the CH in-phase transverse and longitudinal relaxation rates, denoted by *R*(*D*_*y*_) and *R*(*D*_*z*_), respectively. Figure 7A&B compares the cNVT – 2 results for 50 methyls in ubiquitin against experimental data ^28–29^. Overall good agreement is seen, but with overestimation in *R*(*D*_*z*_) for a few residues (e.g., I3 C_γ2_H).

**Figure 7.**
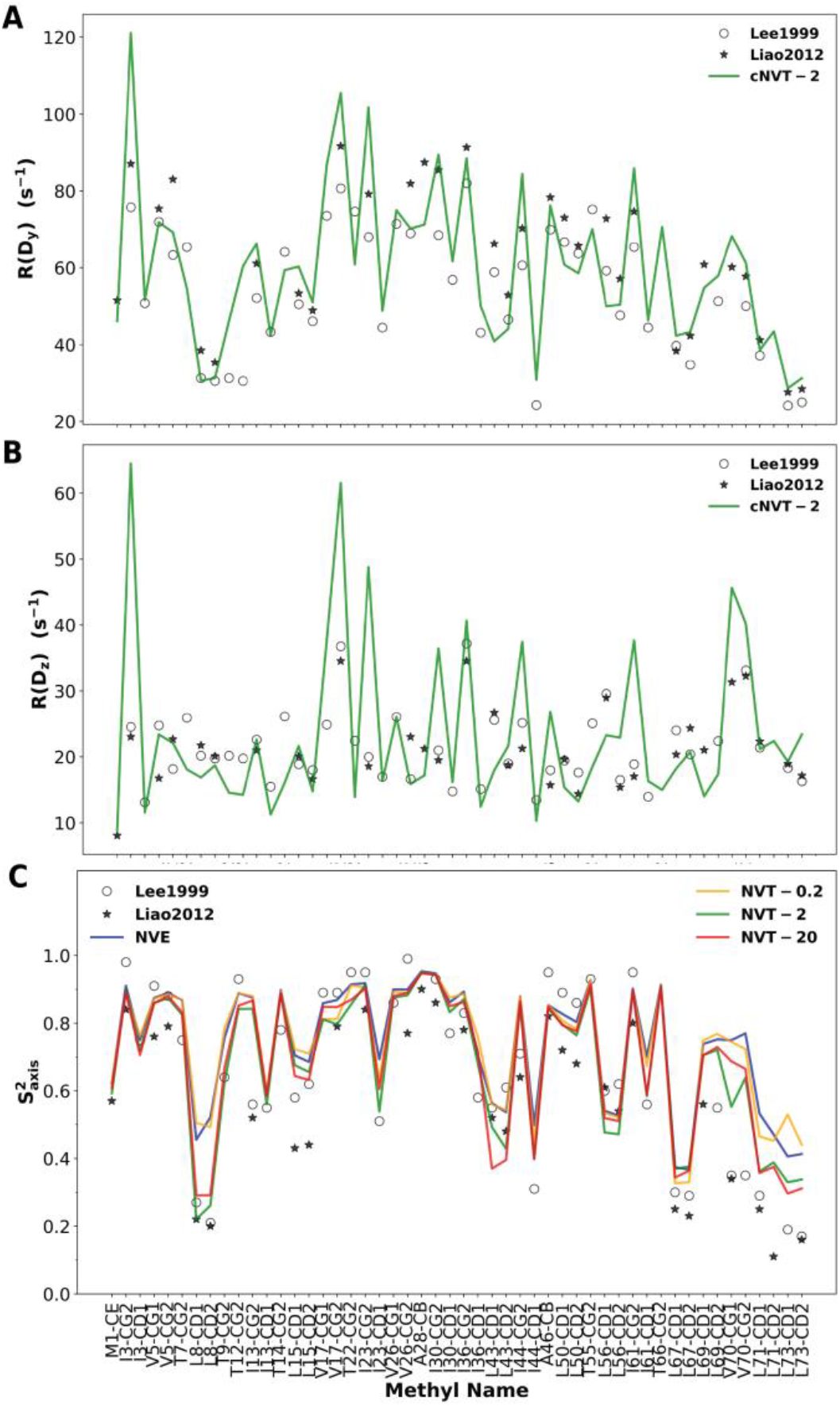
Comparison of side-chain methyl NMR relaxation properties predicted by corrected NVT simulations against experimental data for ubiquitin. (*A*) & (*B*) Side-chain methyl CH in-phase transverse and longitudinal relaxation rates, from corrected NVT (at *ζ* = 2 ps^−1^) and from experimental studies of Lee et al. (28) and Liao et al. (29). (*C*) Methyl axis order parameters, 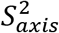, predicted by the *A*_1_ amplitudes of the CC correlation functions in NVE and NVT simulations (at *ζ* = 0.2, 2, and 20 ps^−1^), and measured by Lee et al. and by Liao et al.

As a final validation that the Langevin thermostat does not perturb motional amplitudes, we calculated methyl axis order parameters, 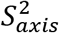, from NVE and NVT simulations. Figure 7C displays 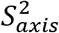 results for 50 methyls in ubiquitin from NVE, NVT – 0.2, NVT – 2, and NVT – 20 as well as from experiments ^28–29^. Overall, 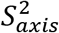 values agree well among the different simulations. However, there is a gap between NVE and NVT – 0.2 values and NVT – 2 and NVT – 20 values for some of the most flexible methyls (indicated by low 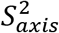), including those on L8, L43, L71, and L73. The NVT – 2 and NVT – 20 values are in closer agreement with experiments, and thus the higher 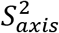 from NVE and NVT – 0.2 simulations is a sign that they suffer from insufficient conformational sampling. The RMSEs of the NVE and NVT – 0.2 simulations from Lee et al. ^28^ are 0.15 and 0.16, respectively. These are already relatively low errors for methyl order parameters. The RMSEs further decrease to 0.11 and 0.12, respectively, for NVT – 2 and NVT – 20 simulations.

## 4. DISCUSSION

We have thoroughly assessed the distortions of the dynamics of globular proteins by the Langevin thermostat, and developed a correction scheme to remove the distortions. The Langevin thermostat dilates the time constants of overall and internal motions, but does not affect the amplitudes of these motions. The extents of dilation are significant for the slower (ns) overall rotation but modest for the fast (sub-ns) internal motions. The correction scheme involves contraction of the time constants, with the contraction factor a linear function of the time constant to be corrected. The corrected dynamics of eight proteins are validated by NMR data for rotational diffusion and for backbone amide and side chain methyl relaxation.

Thermostat distortions observed in limited past studies can now be understood in light of our findings. For example, the sub-ps underdamped time correlation function of a side-chain torsion angle was preserved by a Langevin thermostat at *ζ* = 2 ps^−1^, but became overdamped at *ζ* = 20 and 50 ps^−1 12^. This observation is in line with slight *τ*_2_ dilation at *ζ* = 2 ps^−1^ but clear *τ*_2_ dilation at *ζ* = 20 ps^−1^ in our simulations. Basconi and Shirts ^17^ observed several-fold slowdown in water diffusion and homopolymer chain decorrelation by a Langevin thermostat at *ζ* = 10 ps^−1^. For this damping constant, our Eq [13] predicts a value of 2.8 for the *a* coefficient, meaning that all time constants are increased by at least 2.8-fold by the Langevin thermostat. Debiec et al. ^18^ reported RCTs for GB3, ubiquitin, and binase from NVE and NVT simulations at *ζ* = 1 ps^−1^. For this damping constant, our Eqs [13] and [14] yield *a* = 1.09 and *b* = 0.059 ns^−1^. Applying the resulting time-contraction correction to their NVT results, we predict RCTs of 3.58, 4.49, and 6.34 ns for the three proteins in NVE simulations. These predictions agree well with the reported values, 3.47, 4.62, and 7.26 ns. Lastly Hoffmann et al. ^23^ reported that NPT and NVE simulations produced “very similar” results for methyl axis order parameters and CH in-phase transverse and longitudinal relaxation rates. Their observation on 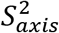 supports our finding that motional amplitudes are not perturbed by thermostats. Also, *R*(*D*_*y*_) and *R*(*D*_*z*_) are dominated by fast (sub-ns) local motions (as *A*_2_ ≫ *A*_1_ for methyl CH bonds) and hence should not be perturbed much either.

We anticipate that our correction scheme for thermostat distortions will have wide applications. Although isotropic overall rotation is modeled in the present study, the correction scheme can be easily extended to model anisotropic rotation of folded proteins or complexes that deviate significantly from a globular shape. Another tantalizing application is to remove dynamic distortions of intrinsically disordered proteins (IDPs) in NVT simulations. As illustration, we apply the correction of Eq [12] to our previous NVT simulations of ChiZ at *ζ* = 3 ps^−1^ ^34^. The correction indeed brings the calculated backbone NH relaxation properties into closer agreement with experimental data (Fig. S10). Most significantly, since the extent of time dilation by a Langevin thermostat grows with increasing timescales of motions, we expect that time dilation for functional important conformational switches ^52^ and protein folding/unfolding that occur on μs or longer timescales will exceed the 2-fold observed here for 5-ns motions. As NVE simulations suffer from temperature drifts ^13^ (Fig. S11) as well as insufficient conformational sampling, it will be impractical to use them for studying IDP dynamics and other slow processes. A thermostat and the necessary dynamic correction provide the only way to propel MD simulations to be on par with NMR and other experimental techniques in determining dynamic properties of proteins.

That the Langevin thermostat does not affect motional amplitudes can be rationalized if we recognize that motional amplitudes are thermodynamic, not dynamic properties, and the Langevin thermostat, by design, preserves thermodynamic properties. That the Langevin thermostat dilates time constants of overall rotation to a much greater extent than those of internal motions can also be rationalized as follows. In overall rotation, a protein experiences only the solvent friction in NVE simulations. A Langevin thermostat not only increases the solvent friction (due to Langevin dynamics of the solvent) but also directly adds friction to each protein atom (due to Langevin dynamics of the protein). The total increase in friction may be comparable in magnitude to the original solvent friction. In contrast, for internal motions, the dominant contribution is internal friction arising from a rugged energy surface, and hence the increase in friction by the Langevin thermostat is less important. It is less straightforward to explain why time dilation for overall rotation grows with RCT and hence protein size (Fig. S1). The RCT from an NVE simulation is proportional to the original solvent viscosity *η*_0_ and the protein volume *V*: RCT_NVE_ = *βη*_0_*V*, where *β* is a constant. Correspondingly we have RCT_NVT_ = *β*(*η*_0_ + *η*_1_)*V*, where *η*_1_ is the total increase in viscosity by the Langevin thermostat. If *η*_1_ depends on protein size, then the time dilation, RCT_NVT_/RCT_NVE_, will also depend on protein size. These foregoing qualitative analyses are also instructive for other types of thermostats.

Just as water models affect the RCT of a protein ^13, 20^, we should expect that the dynamic distortions of a Langevin thermostat depend on the water model. As already noted, the dynamic distortions also depend the timescales of motions and on protein size. Without corrections, variations in distortions due to these inter-related factors can greatly complicate force-field parameterization and validation and affect the transferability of force fields. Our correction scheme will be valuable for both force-field parameterization and transferability.

## ASSOCIATED CONTENT

### Supporting Information

The Supporting Information is available free of charge. Two tables listing rotational correlation times and root-mean-square-errors and 11 figures displaying time dilation by a Langevin thermostat; backbone NH and side-chain methyl CH time correlation functions; parameters of bi-exponential fits for NH time correlation functions; dependences of *τ*_1_ and *τ*_2_ on the damping constant of the Langevin thermostat; comparison of backbone NH NMR relaxation properties predicted by NVE and NVT simulations against experimental data; backbone NH order parameters; parameters of bi-exponential fits for ubiquitin methyl CH time correlation functions; comparison of backbone NH NMR relaxation properties calculated from NVT simulations of ChiZ against experimental data; and temperature drift in NVE simulations and temperature regulation by a Langevin thermostat (PDF)

## AUTHOR INFORMATION

### Author Contributions

A.H. and H.-X.Z. designed the research. A.H. and M.M. performed the research and analyzed the data. A.H. and H.-X.Z. wrote the manuscript.

### Notes

The authors declare no competing financial interests.

## ACKNOWLEDGMENT

This work was supported by National Institutes of Health Grant R35 GM118091.

## ABBREVIATIONS

BPTI: bovine pancreatic trypsin inhibitor
GB3: the third immunoglobulin-binding domain of protein G
HEWL: hen egg white lysozyme
MD: molecular dynamics
NVE: particle number / volume / energy
NVT: particle number / volume / temperature
PME: particle mesh Ewald
RCT: rotational correlation time
RMSE: root-mean-square-error
RNase: ribonuclease
SASA: solvent accessible surface area

