## Supporting Tables and Figures for "Removing Thermostat Distortions of Protein Dynamics in Constant-Temperature Molecular Dynamics Simulations"

*Supporting Information*

**Table S1** Rotational correlation times (RCTs) of eight proteins from experimental studies, from NVE simulations, and from NVT simulations at three damping constants (0.2, 2, 20 $ps^{-1}$)

| Protein | Temperature (°K) | RCT (ns) | | | | |
|---|---|---|---|---|---|---|
| | | EXP | NVE | NVT-0.2 | NVT-2 | NVT-20 |
| θ-Defensin | 298 | 1.09 | 1.21±0.06 | 1.24±0.03 | 1.77±0.05 | 5.74±0.14 |
| Trp-Cage | 300 | 1.1 | 0.96±0.02 | 0.99±0.02 | 1.43±0.03 | 5.06±0.02 |
| ShK | 283 | 2.6 | 3.39±0.32 | 3.36±0.04 | 5.06±0.14 | 18.75±0.77 |
| GB3 | 300 | 3.43±0.03 | 3.22±0.12 | 3.51±0.03 | 6.09±0.12 | 26.03±0.37 |
| BPTI | 298 | 3.5 | 3.92±0.48 | 3.96±0.15 | 6.82±0.11 | 27.98±0.15 |
| Ubiquitin | 300 | 4.1 | 4.33±0.09 | 4.92±0.13 | 8.77±0.17 | 39.83±1.73 |
| RNase T1 | 308 | 5.30±0.13 | 4.73±0.12 | 5.43±0.17 | 10.50±0.55 | 46.98±1.93 |
| HEWL | 308 | 5.7±0.2 | 5.79±0.39 | 6.50±0.22 | 13.42±0.37 | 66.63±1.07 |

**Table S2** Root-mean-square-errors (RMSEs) of calculated NH longitudinal and transverse relaxation rates and heteronuclear Overhauser effects

| | θ-Defensin | Trp-Cage | ShK | GB3 | BPTI | Ubiquitin | RNase T1 | HEWL |
|---|---|---|---|---|---|---|---|---|
| $R_1$ RMSE ($s^{-1}$) | | | | | | | | |
| NVE | 0.12 | 0.53 | 0.16 | 0.18 | 0.16 | 0.17 | 0.22 | 0.36 |
| cNVT-0.2 | 0.12 | 0.54 | 0.12 | 0.16 | 0.17 | 0.14 | 0.18 | 0.36 |
| cNVT-2 | 0.11 | 0.49 | 0.13 | 0.15 | 0.16 | 0.14 | 0.18 | 0.35 |
| cNVT-20 | 0.13 | 0.49 | 0.11 | 0.12 | 0.18 | 0.14 | 0.24 | 0.33 |
| $R_2$ RMSE ($s^{-1}$) | | | | | | | | |
| NVE | -- | 1.2 | 1.2 | 0.33 | 0.96 | 0.55 | 1.5 | 0.63 |
| cNVT-0.2 | -- | 1.2 | 1.2 | 0.36 | 0.92 | 0.60 | 1.3 | 0.55 |
| cNVT-2 | -- | 1.1 | 1.1 | 0.39 | 0.94 | 0.62 | 1.3 | 0.55 |
| cNVT-20 | -- | 1.1 | 1.2 | 0.56 | 0.96 | 0.69 | 1.5 | 0.63 |
| $^{1}H$-$^{15}N$ Het. NOE RMSE | | | | | | | | |
| NVE | 0.15 | 0.18 | 0.05 | 0.04 | 0.12 | 0.10 | 0.03 | 0.09 |
| cNVT-0.2 | 0.12 | 0.20 | 0.06 | 0.04 | 0.12 | 0.09 | 0.03 | 0.08 |
| cNVT-2 | 0.13 | 0.26 | 0.05 | 0.05 | 0.12 | 0.08 | 0.05 | 0.08 |
| cNVT-20 | 0.19 | 0.25 | 0.05 | 0.05 | 0.10 | 0.20 | 0.06 | 0.09 |

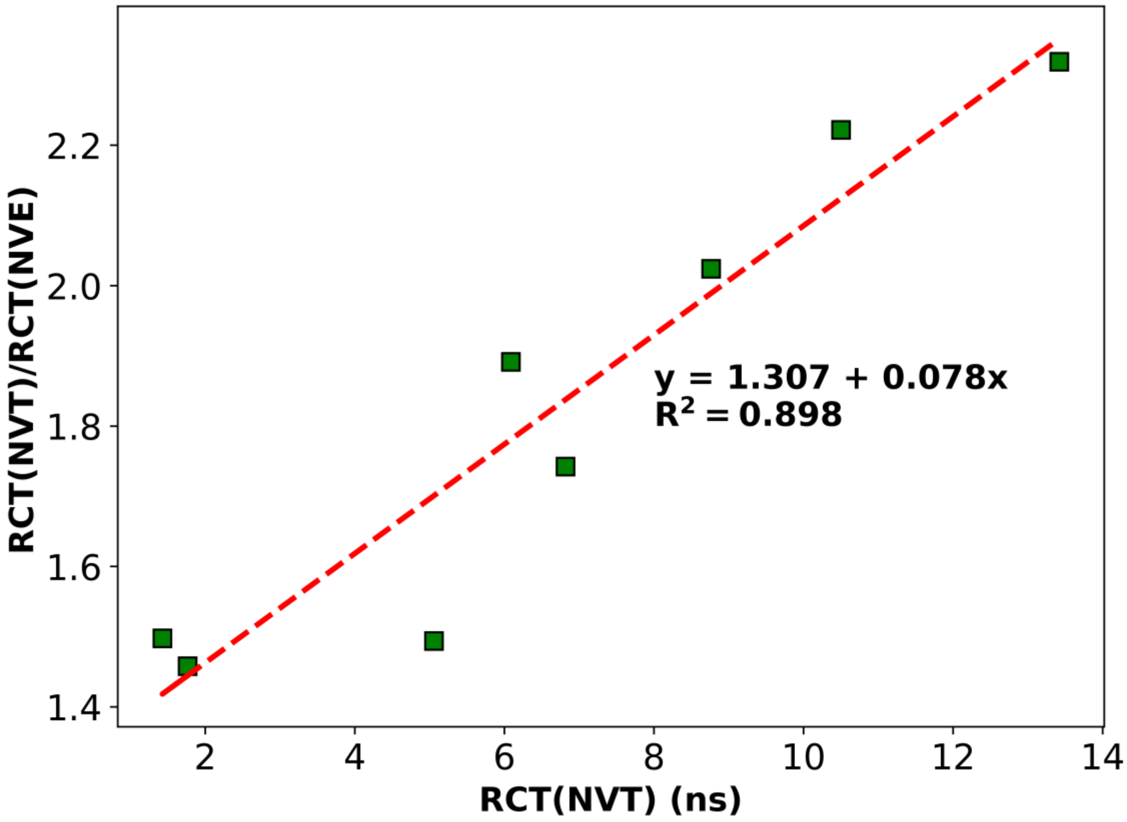


**Figure S1.** The time dilation by a Langevin thermostat grows with protein size. Time dilation is measured by the ratio between the rotational correlation times (RCTs) calculated from NVT and NVE simulations; protein size is measured by RCT. Each data point is for one of the eight proteins in Fig. 1. NVT simulations were regulated by a Langevin thermostat at $\zeta$ = 2 $ps^{-1}$.

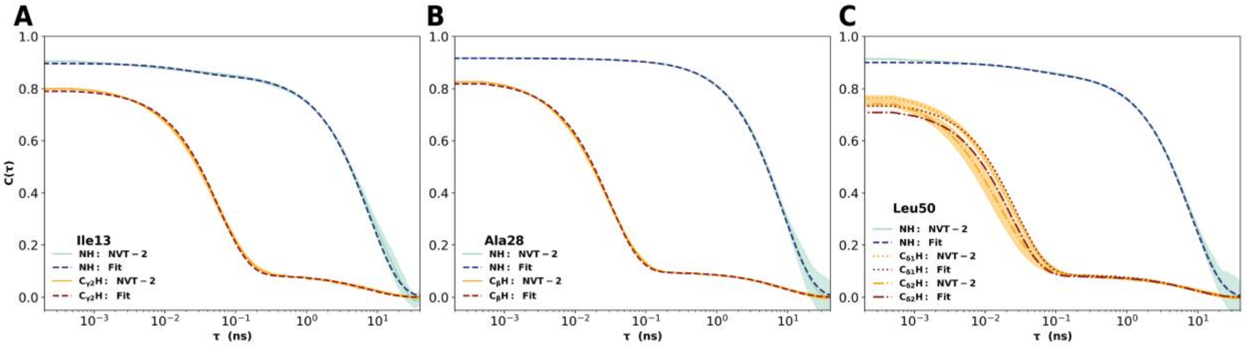


**Figure S2.** Backbone NH and side-chain methyl CH time correlation functions, and their bi-exponential fits. Results are from NVT – 2 simulations. Three residues from ubiquitin are chosen for illustration. The standard deviations among four replicate simulations are shown as shading.

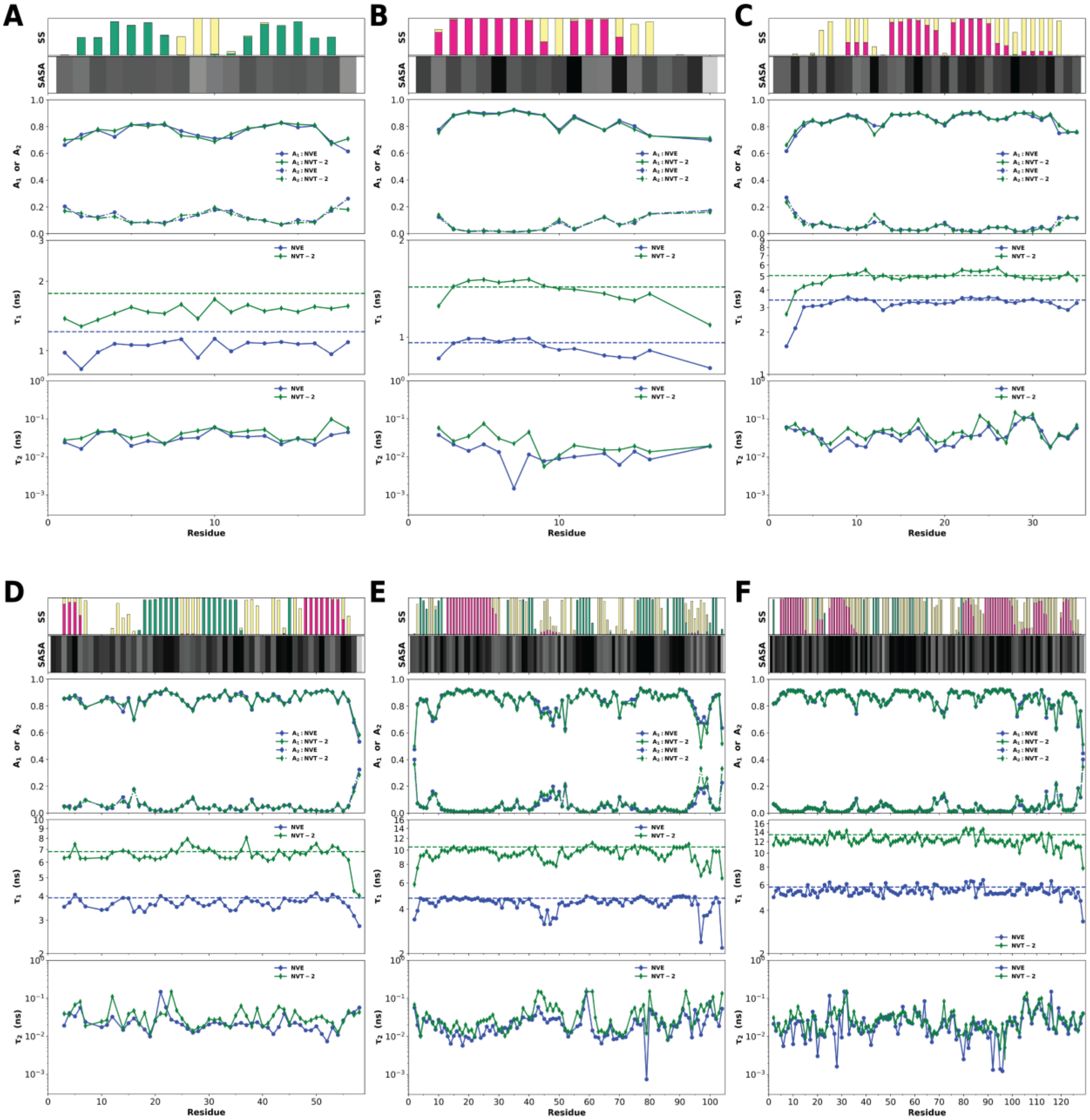


**Figure S3.** Extension of Fig. 3 to six other proteins. (*A*) θ-Defensin. (*B*) Trp-cage. (*C*) ShK. (*D*) BPTI. (*E*) RNase T1. (*F*) HEWL. In each panel, from top to bottom, the following items are displayed: secondary structure (SS); percent of solvent exposure; $A_1$ and $A_2$ amplitudes of the bi-exponential fits; $\tau_1$ time constant, with rotational correlation times displayed as dashed horizontal lines; $\tau_2$ time constant.

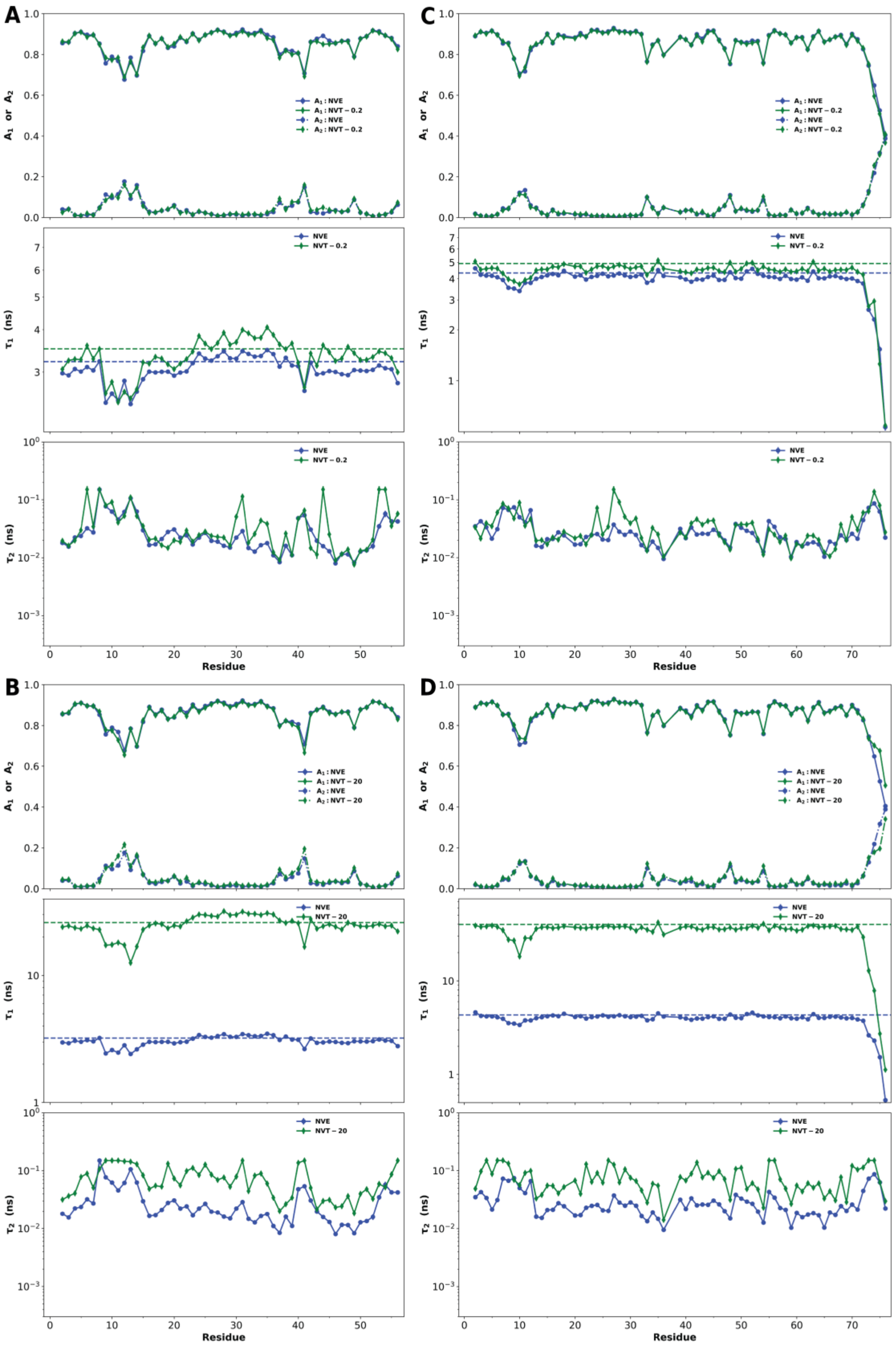


**Figure S4.** Extension of Fig. 3 to NVT simulations at $\zeta$ = 0.2 and 20 $ps^{-1}$. (*A*&*B*) GB3. (*C*&*D*) Ubiquitin.

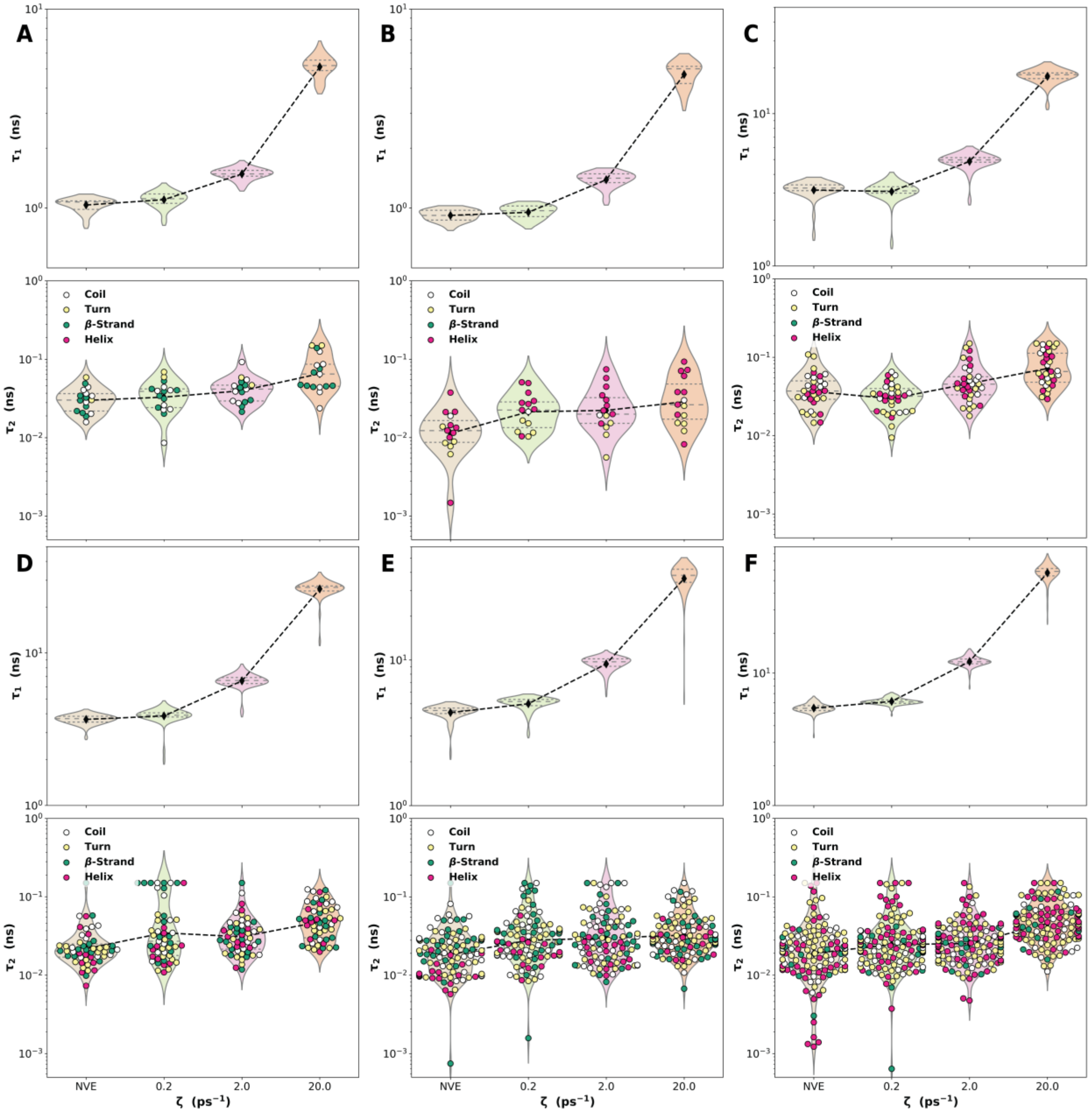


**Figure S5.** Extension of Fig. 4 to six other proteins. (*A*) θ-Defensin. (*B*) Trp-cage. (*C*) ShK. (*D*) BPTI. (*E*) RNase T1. (*F*) HEWL.

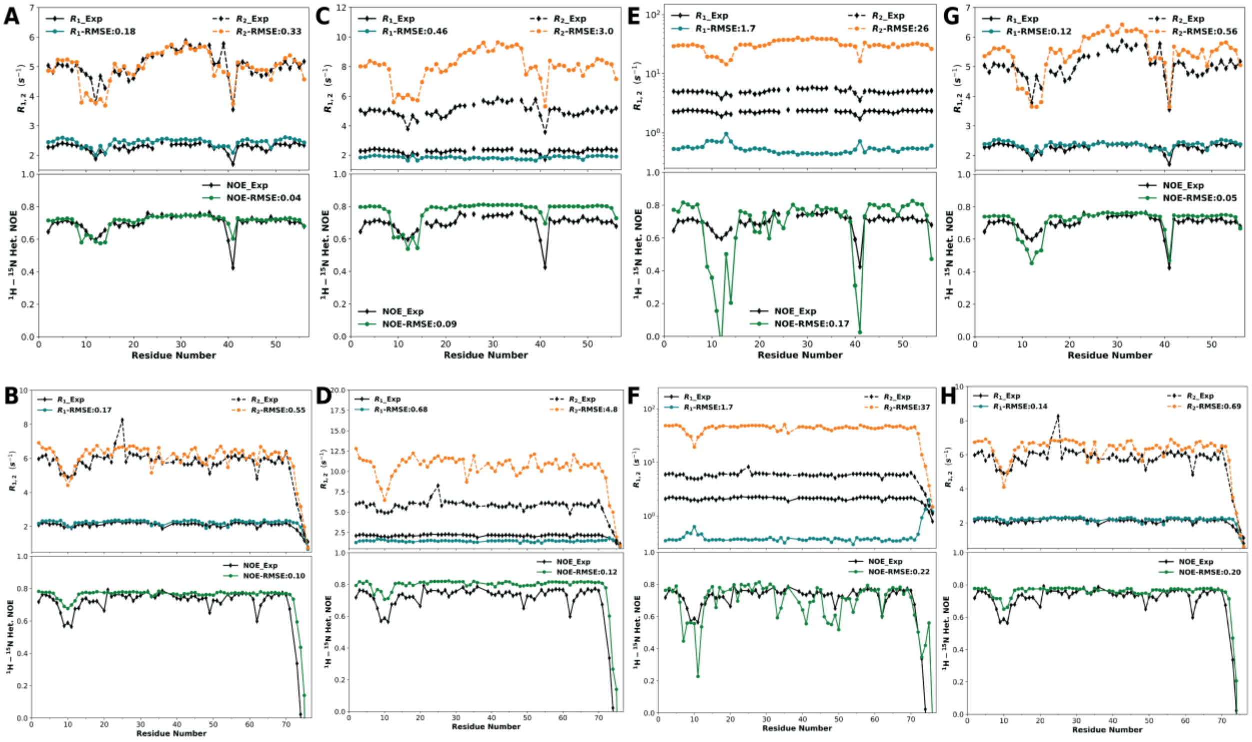


**Figure S6.** Comparison of backbone NH NMR relaxation properties predicted by NVE and NVT simulations against experimental data. (*A*), (*C*), (*E*), & (*G*) are for GB3 and (*B*), (*D*), (*F*), & (*H*) are for ubiquitin. (*A*) & (*B*) NVE results. (*C*) & (*D*) NVT results at $\zeta$ = 2 $ps^{-1}$. (*E*) & (*F*) NVT results at $\zeta$ = 20 $ps^{-1}$. (*G*) & (*H*) Corrected NVT results at $\zeta$ = 20 $ps^{-1}$. In each panel, results from MD simulations are shown in color, while experimental data are shown in black.

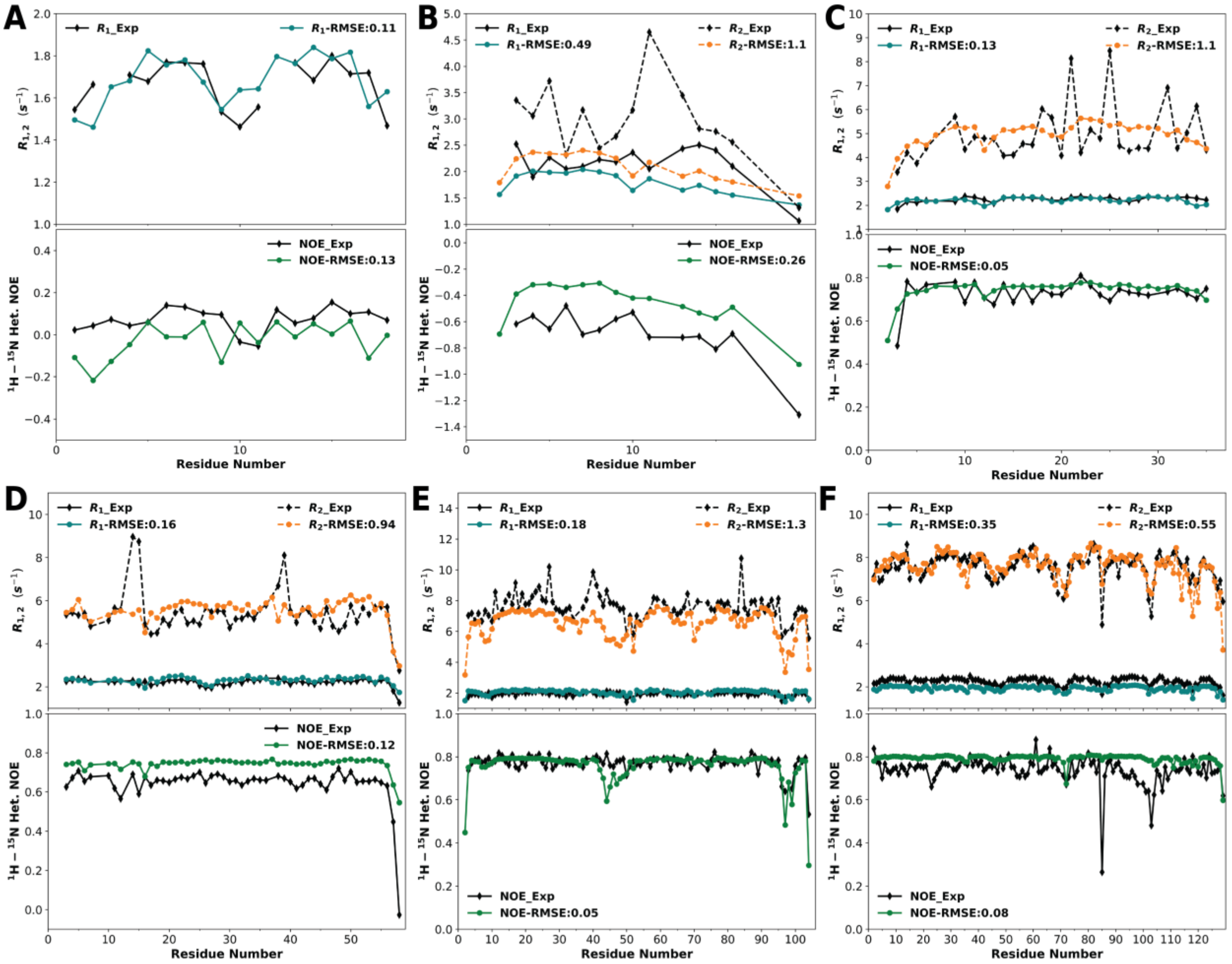


**Figure S7.** Extension of Fig. 6 to six other proteins. (*A*) θ-Defensin. (*B*) Trp-cage. (*C*) ShK. (*D*) BPTI. (*E*) RNase T1. (*F*) HEWL.

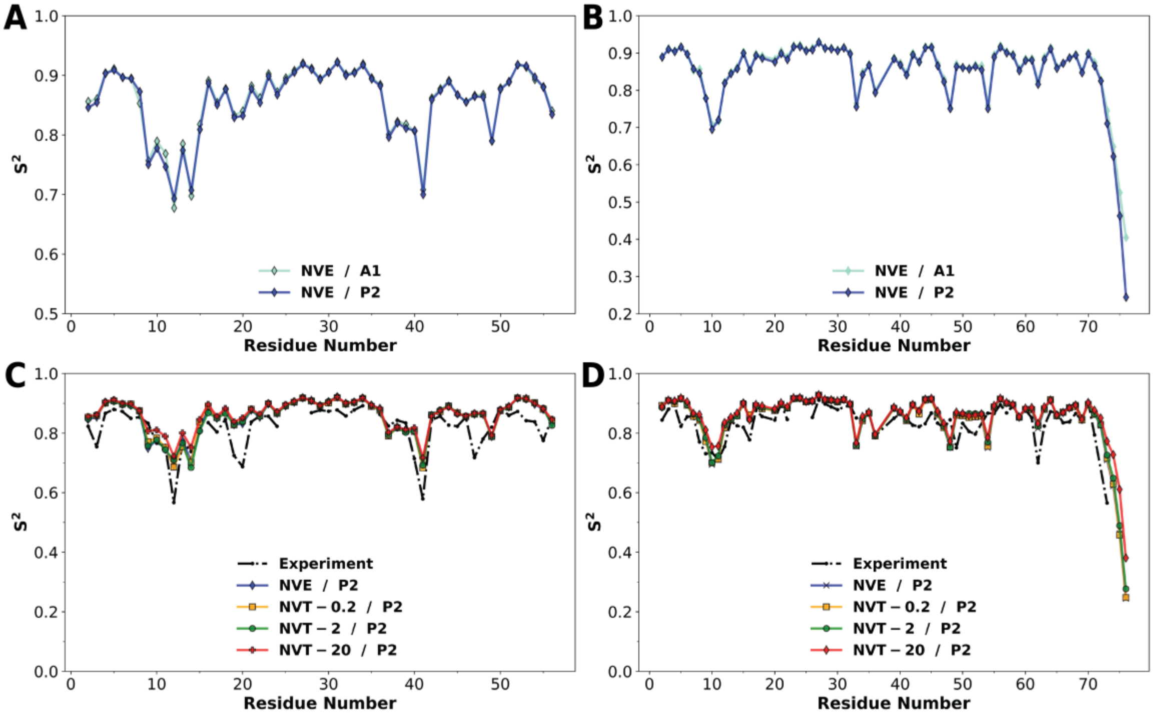


**Figure S8.** Backbone NH order parameters, $S^2$. (*A*) & (*C*) are for GB3 and (*B*) & (*D*) are for ubiquitin. (*A*) & (*B*) Comparison of order parameters from the A1 and P2 methods. (*C*) & (*D*) Comparison of order parameters calculated by the P2 method among NVE, NVT – 0.2, NVT – 2, and NVT – 20 simulations, and also against experimental data. The P2 method used 1-ns blocks for structural alignment.

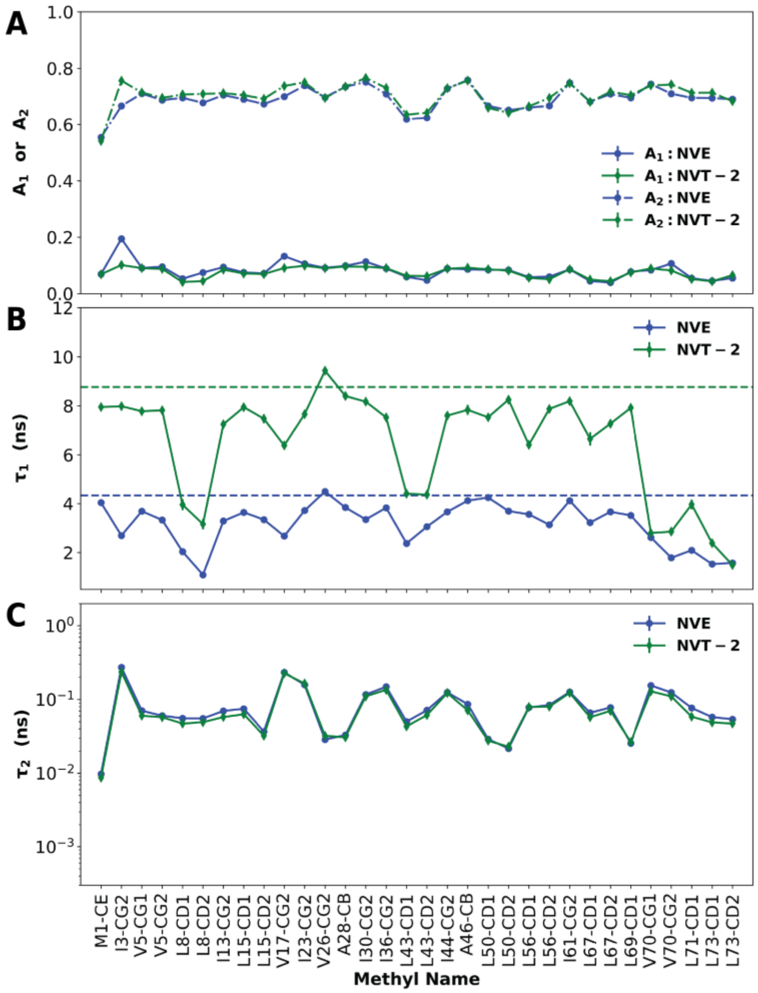


**Figure S9.** Parameters of bi-exponential fits for ubiquitin methyl CH time correlation functions. Results for NVE and NVT (at $\zeta$ = 2 $ps^{-1}$) simulations are shown. (*A*) Amplitudes $A_1$ and $A_2$ of the bi-exponential fits. (*B*) Time constant $\tau_1$. The rotational correlation times calculated from these simulations are displayed as dashed horizontal lines. (*C*) Time constant $\tau_2$.

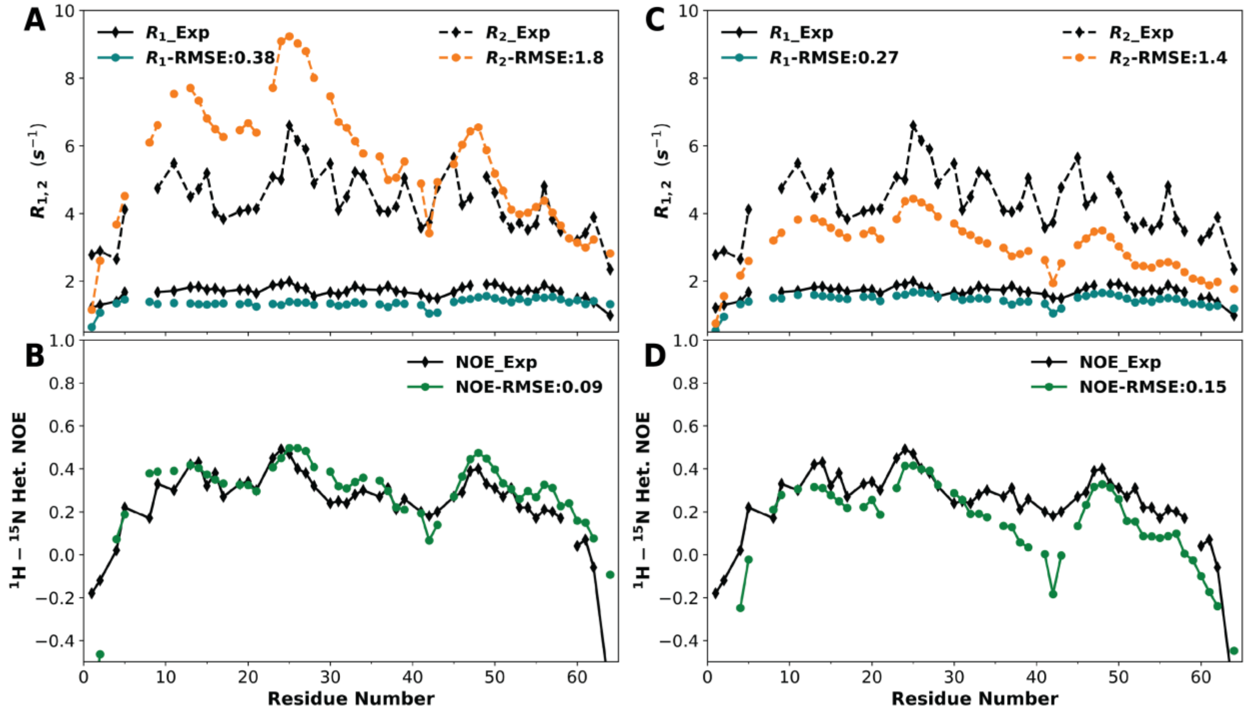


**Figure S10.** Comparison of backbone NH NMR relaxation properties calculated from NVT simulations of ChiZ against experimental data. (*A*) & (*B*) Results directly from NVT (at $\zeta$ = 3 ps$^{-1}$) simulations shown in color; experimental data shown in black. (*C*) & (*D*) The same comparison but for simulation results after time-contraction correction.

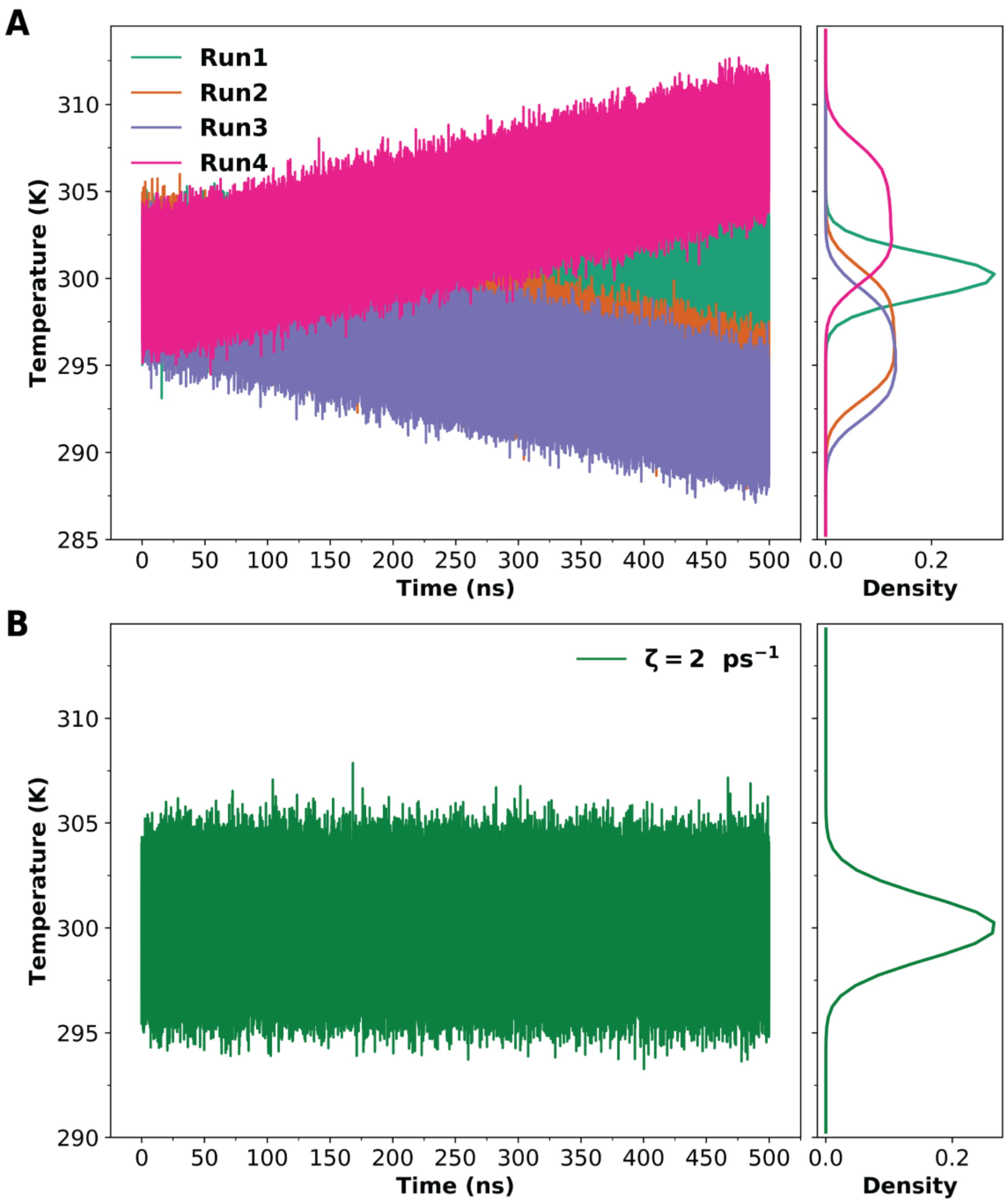


**Figure S11.** Temperature drift in NVE simulations and temperature regulation by a Langevin thermostat at $\zeta = 2$ ps$^{-1}$. (*A*) Time traces of temperature in four replicate NVE runs. Temperature is stable in run1, but drifts upward in run4 and downward in run2 and run3. (*B*) A representative time trace of temperature in a Langevin simulation, showing stable temperature.
